# Natural selection acts on epigenetic marks

**DOI:** 10.1101/2020.07.04.187880

**Authors:** Leandros Boukas, Afrooz Razi, Hans T. Bjornsson, Kasper D. Hansen

## Abstract

In the study of molecular features like epigenetic marks, it is appealing to ascribe observed patterns to the action of natural selection. However, this conclusion requires a test for selection based on a well-defined notion of neutrality. Here, focusing on epigenetic marks at gene loci, we formalize what it means for an epigenetic mark to be neutral, and develop a test for selection. Our test respects the foundational aspect of epigenetics: *trans*-regulation by transcription factors and chromatin modifiers. It also enables adjustment for confounders. We establish that promoter DNA methylation, promoter H3K4me3 and exonic H3K36me3 are all under selection, and that this is unlikely to be a passive consequence of selection on gene expression. The effect of these epigenetic marks on fitness is arguably partly explained by a causal involvement in gene regulation. However, we also investigate the complementary explanation that DNA methylation and H3K36me3 are under selection in part because they modify the mutation rate of important genomic regions. We show that this explanation is consistent with empirical observations as well as population genetics theory, because of the *trans*-regulation. Exemplifying the protection of important regions from high mutability, we demonstrate that in humans the more intolerant to loss-of-function mutations a gene is, the lower its coding mutation rate is. Our framework for selection inference is simple but general, and we speculate that its core ideas will be useful for additional molecular features beyond epigenetic marks.

## Introduction

Natural selection is one of the fundamental forces shaping the evolution of living organisms (Fisher, 2015; Crow, Kimura, 2009). As a result, detecting the action of selection is one of the major goals of evolutionary biology. At the molecular level, the focus thus far has been on identifying signatures of selection within the DNA sequence (Bamshad, Wooding, 2003; Nielsen, 2005; Nielsen et al., 2007). Major insights into the action of selection have been attained by comparative genomic analyses, as well as analyses of population-level genetic variation (see Nielsen et al. (2005), Sabeti et al. (2006), Blekhman et al. (2008), Lek et al. (2016), and Cassa et al. (2017) for examples of results in humans).

However, natural selection acts on phenotypes; footprints of selection in the DNA sequence only reflect the extent to which the DNA causally determines these phenotypes. In turn, the causal effect of the DNA sequence on phenotypes is mediated via its effect on intermediate functional molecular features. There is a wide range of features potentially acting as such mediators: from epigenetic marks to the structure and activity of gene regulatory networks and signaling pathways. Therefore, it is of interest to move beyond the DNA sequence, and identify selective pressures on specific molecular features. This is akin to the major objective of “evolutionary cell biology” (Lynch et al., 2014; Lynch, Trickovic, 2020).

In theory, one way to detect selection on a given molecular feature would be to identify the parts of the DNA sequence which determine that feature, and then test for selection on these sequence parts with established methods. However, this approach can be difficult to apply in practice. For example, the genome-wide landscape of epigenetic marks is determined by networks of upstream regulatory factors (transcription factors and chromatin modifiers; Stadler et al. (2011), Krebs et al. (2014), Boulard et al. (2015), and Lappalainen, Greally (2017)); at most genomic locations, the identities of the relevant factors and the logic governing their interactions are not fully understood. Additionally, even if the causally relevant sequence parts are known for a given feature, interpreting signatures of selection on them can be complicated by issues such as pleiotropy (e.g. a given protein may participate in more than one gene regulatory network or signaling pathway, each of them serving a different function).

An alternative approach which overcomes these issues is to directly infer selection on the intermediate feature of interest. The major barrier to achieving this is the lack of a suitable concept of neutrality. For instance, it may be tempting to attribute the distribution of a certain epigenetic mark across the genome to the action of selection. But how would that same mark be distributed if it were neutral? Without a neutrality concept, there is no null hypothesis against which the alternative of selection can be tested (Lynch, 2007; Lynch, Trickovic, 2020; Eres, Gilad, 2021).

Hereafter, we restrict our focus to epigenetic marks. We propose that recently derived estimates of selection coefficients of coding loss-of-function alleles allow for a natural definition of the fitness effect of epigenetic marks associated with genes, and we show that this definition can be used to derive a test for selection from first principles. We then apply this test to several marks and explore the biological basis and implications of the results.

## Results

### An approach for inferring selection on epigenetic marks

In what follows, we refer to epigenetic marks with different possible “epigenetic states”. For instance, the mark can be DNA methylation at proximal promoters, and in that case each promoter can have either the methylated or the hypomethylated state (details for why this binary labeling is appropriate provided in next section). Our general goal for any given mark is to infer whether a particular state is favored, or acted against, by natural selection.

A key aspect of the biology of epigenetic marks, as mentioned above, is that their presence is controlled simultaneously at multiple locations, via the action of *trans*-acting regulators: transcription factors and chromatin modifiers. Experimental evidence suggests that different genomic locations are under the control of different combinations of *trans*-acting factors, which can interact in different ways (Stadler et al., 2011; Krebs et al., 2014; Boulard et al., 2015). To capture this, we associate each epigenetic mark with its “trans-regulatory groups”. Each *trans*-regulatory group is a set of locations which are all regulated by the same factors, with the same regulatory logic defining how these factors interact. Therefore, all locations in a trans-regulatory group have the same state of the epigenetic mark. Figure 1a depicts a simple hypothetical example with a binary epigenetic mark and two regulatory groups, resulting in 4 possible states (see also Supplemental Figure S1 for examples of different potential regulatory logics). Naturally, between-individual variation in the expression and activity of the upstream factors controlling a trans-regulatory group can result in between-individual variation in the epigenetic state of that group (Figure 1a). Hereafter, we restrict our attention to locations associated with genes (e.g. promoters).

**Figure 1.**
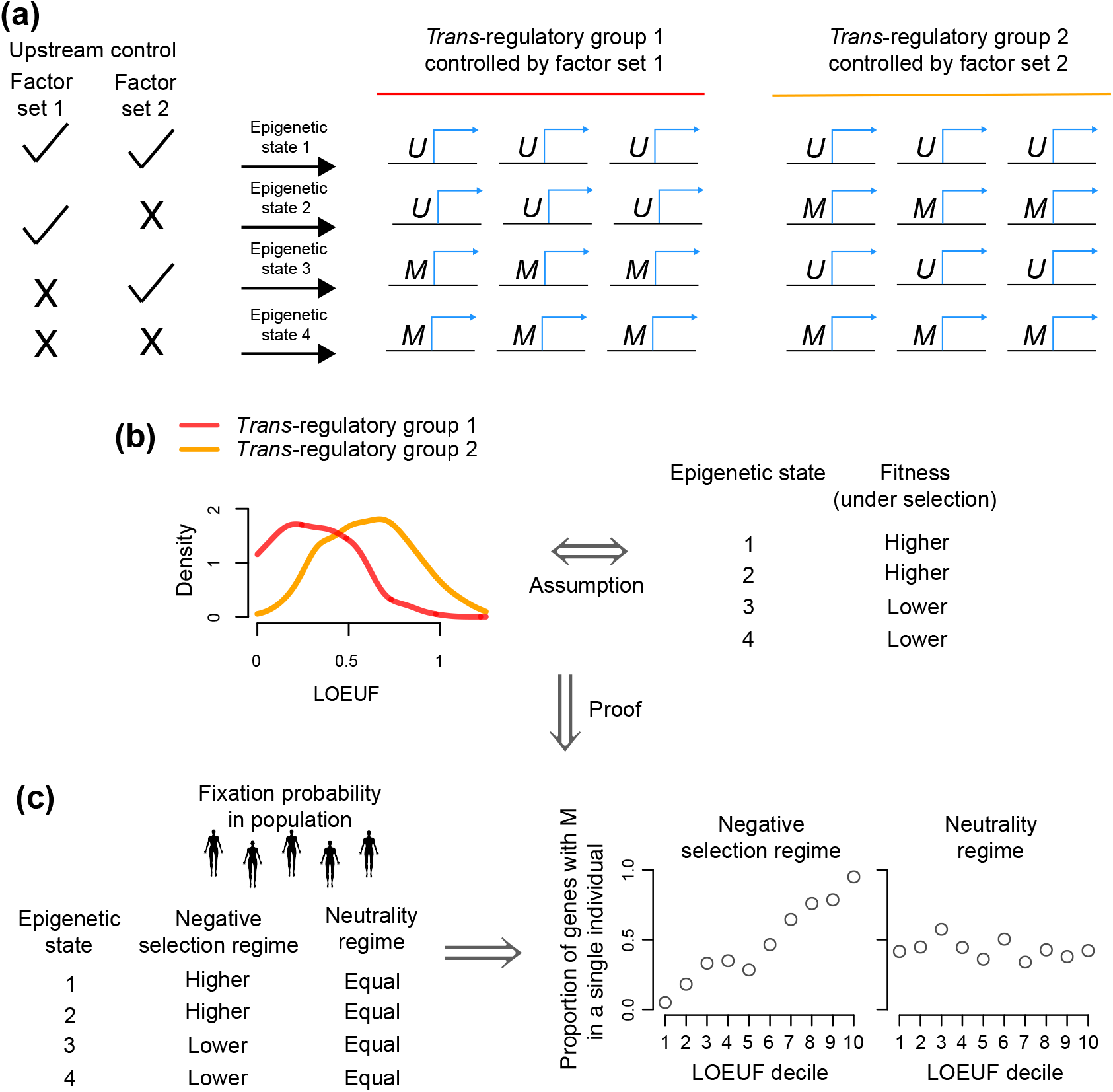
Intuition behind our test for selection on epigenetic marks. We consider a hypothetical example with an epigenetic mark that has two possible states (*U, M*). **(a)** Two *trans*-regulatory groups controlled by two different sets of upstream factors. When the set of upstream factors are active, each promoter in the corresponding group has the *U* state. **(b)** An illustration of the assumption of a relationship between the distribution of selection coefficients against heterozygous loss-of-function alleles (quantified by LOEUF; a proxy for *s*_het_) of the genes in each *trans*-regulatory group, and the associated fitness of individuals with different epigenetic states when the mark is under selection (see Methods for details). **(c)** Under assumption (b), we prove (Methods) that different evolutionary regimes give rise to a different kind of relationship between the fixation probability of a given epigenetic state and individual LOEUFs, even when the *trans*-regulatory groups are unknown. This manifests as a relationship between the probability of observing the epigenetic state at a given gene and its LOEUF, and can be visualized across groups of genes, when genes are ranked based on LOUEF (e.g. deciles).

Each *trans*-regulatory group has an associated fitness *w_G_*, which is the fitness of individuals with the epigenetic state of interest at that group. This fitness represents the aggregation of the fitness effects of that state at each individual gene belonging to the group; the state does not need to have the same fitness effect at each of these genes. Our key modeling step is to assume that the group-level fitness *w_G_* is a function of the individual *s*_het_ coefficients of the genes in the group (Methods), where *s*_het,*g*_ is the selection coefficient of a heterozygous coding loss-of-function allele at a given gene *g*. In other words, we assume that the mark would have a different fitness effect at the group if the corresponding *s*_het_ coefficients were different. For illustration purposes, we show how *w_G_* and the genic *s*_het_’s are related under the additional assumption of a multiplicative fitness landscape (Methods), though we note that our inference procedure makes no such assumption. Because each *trans*-regulatory group has its own *w_G_*, the epigenetic mark can be under selection at some groups but not under selection at others. We say the epigenetic mark is under selection if it is under selection at one or more group(s).

To perform inference about selection, the major practical challenge is that the *trans*-regulatory groups are not well-characterized. The reason is that it is not known precisely which transcription factors and chromatin modifiers control the epigenetic state at most locations in the genome, and precisely how this control is achieved; recent work on how Polycomb Repressive Complex 2 affects promoter DNA methylation is illustrative of this complexity (Methods; Boulard et al. (2015)). Despite this unknown regulatory structure, we show (Methods) that it is possible to derive a statistical test that can distinguish between different evolutionary scenarios with regards to the fitness effect of an epigenetic mark: neutrality, negative selection, and positive selection. Our test relies on the assumption that differences in the fitness effect of a given epigenetic mark at two different trans-regulatory groups is associated with differences in the shet distributions of the genes in the two groups (Figure 1b; Supplemental Figure S2), and is based on a quantity we term dM. This quantity captures the relationship between the fixation probability of an epigenetic mark and genic *s*_het_’s across the genome (Methods). Under neutrality, *dM* = 0, meaning that the epigenetic mark has the same fixation probability independently of *s*_het_, and thus is equally likely to be observed across genes (Figure 1c). By contrast, under negative selection, the epigenetic mark is less likely to be fixed at genes with high *s*_het_, yielding *dM* < 0 and a (stochastically) monotonic relationship between the presence of the mark and *s*_het_ across the genome (Figure 1c); the opposite is true under positive selection (Methods).

It is worth noting that our approach leverages the fact that a given epigenetic mark occurs multiple times throughout the genome (e.g. DNA methylation or H3K4me3 are encountered at many promoters). Because of this, the test can be applied using only a single genome; it does not require population-scale data or data across species. Implicitly, this is based on the assumption that selection acts in the same direction (positive or negative) at each gene where it is present (Methods). Later in the text, we provide specific examples which argue that gene-specific optimization of epigenetic states via negative selection at some genes, and positive selection at others, is unlikely.

Finally, while we have framed the approach using *s*_het_ coefficients, in practice any metric that reflects the selective effects of heterozygous coding loss-of-function alleles can be used. Several such metrics have been developed recently, from large-scale analyses of exome sequences (Petrovski et al., 2013; Lek et al., 2016; Cassa et al., 2017; Karczewski et al., 2020). For our analyses here we use LOEUF from gnomAD (Karczewski et al., 2020), with smaller LOEUF values indicating greater selective pressure (Karczewski et al., 2020). We verified that there is strong concordance between LOEUF and *s*_het_ estimates (Supplementary Figure S3b; Spearman’s *ρ* = −0.85), which ensures that the results are robust to this choice.

### Testing for selection on different epigenetic marks

We applied our test to investigate the selective pressure on different epigenetic marks. To illustrate the generality of our approach, we considered 3 different marks (DNA methylation, H3K4me3, H3K36me3) and a broad range of genomic compartments (promoters, gene bodies, transcriptional end regions).

### Proximal promoter DNA methylation in the male germline is under negative selection

We first focused on promoter DNA methylation in the male germline. We started by exploring the pattern of methylation variation across promoters, using whole-genome bisulfite sequencing data from human sperm (Methods). We observed that proximal promoters (defined here as +/− 500bp from the TSS), are either completely methylated or completely hypomethylated (Supplemental Figure S4a). Therefore, we treated proximal promoter methylation as a binary mark.

We investigated the distribution of proximal promoter methylation with respect to downstream gene loss-of-function intolerance, and found a strong relationship: greater loss-of-function intolerance is strongly associated with a smaller probability of proximal promoter methylation (Figure 2a). At the most loss-of-function-intolerant genes (bottom 10% LOEUF), methylation is almost universally absent, with only 0.8% of proximal promoters being methylated. Our test yields a statistically significant *dM* value of −0.07 (*p* = 9.9 · 10^-5^, Figure 2b), indicating negative selection against proximal promoter methylation.

**Figure 2.**
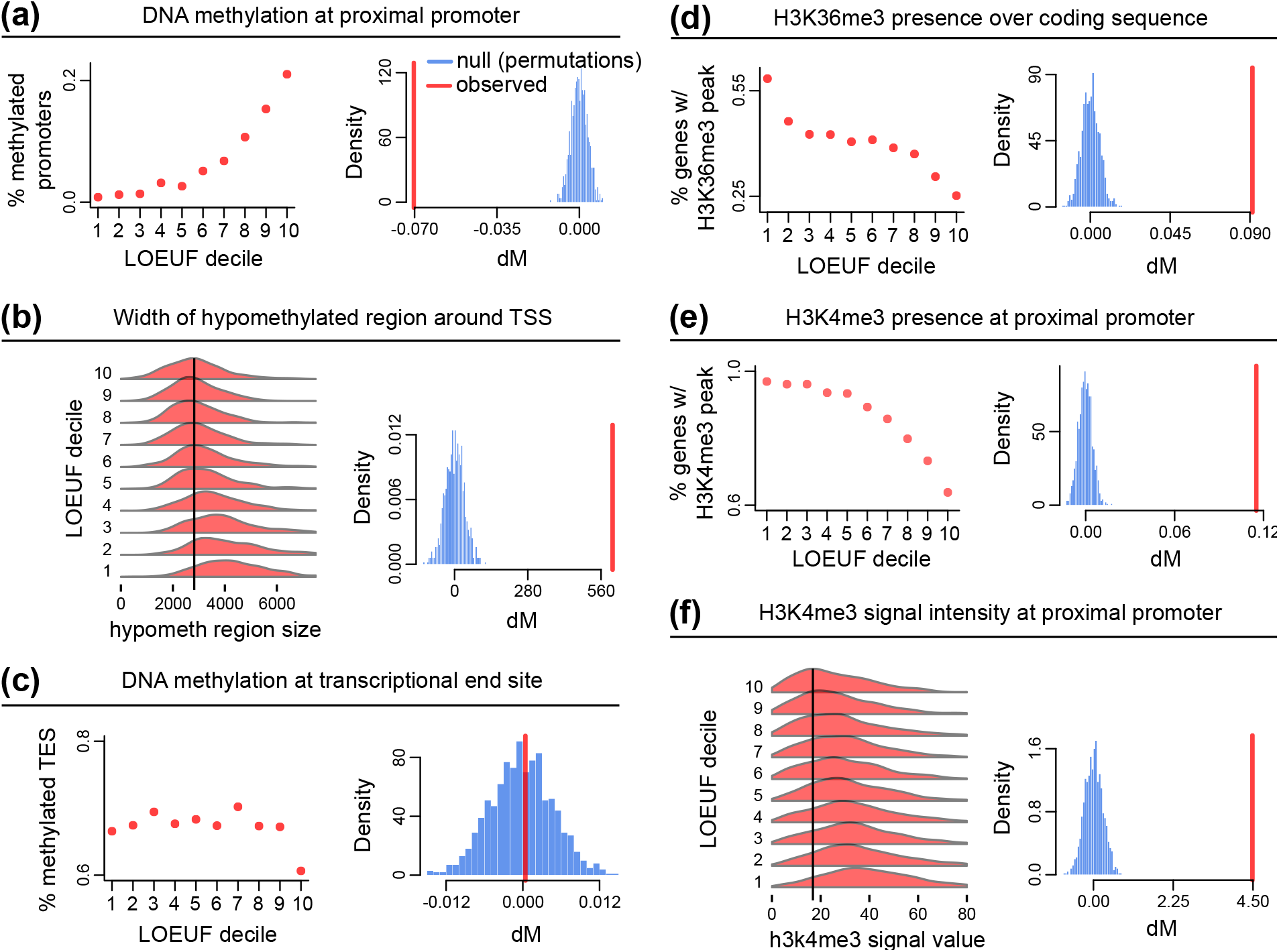
Testing for selection on different epigenetic marks. Each panel consists of two plots. The left plot is a visual depiction of the relationship between the state of the mark and genic loss-of-function intolerance (quantified by LOEUF). The right plot is the observed value of *dM* together with a null distribution (see Methods). **(a)** DNA methylation at the proximal promoter (+/− 500bp around the transcriptional start site; TSS) in the male germline (sperm). **(b)** The width of the hypomethylated region around the TSS at genes with a hypomethylated proximal promoter in the male germline. **(c)** DNA methylation at the transcriptional end region (+/− 500bp) in the male germline. **(d)** Presence of H3K36me3 at the coding sequence in the male germline (sperm). **(e)** Presence of H3K4me3 at the proximal promoter in human H1 embryonic stem cells. **(f)** The ChIP-seq signal intensity of H3K4me3 at genes with H3K4me3 presence at the proximal promoter in H1 embryonic stem cells.

### The width of the hypomethylated region around hypomethylated proximal promoters in the male germline is under directional selection

While each proximal promoter is either completely methylated or hypomethylated, the width of the hypomethylated region around hypomethylated proximal promoters exhibits continuous variation along the genome (Supplemental Figure S4b; Methods). Thus far, such variation has not been shown to have a functional role, unlike the binary methylation state of proximal promoters which has been linked to gene expression regulation. Our approach for selection inference extends to epigenetic marks whose possible states range within a continuous spectrum (Methods), enabling us to examine whether the width of these hypomethylated regions has a functional role without a prespecified hypothesis about what that role may be. We discovered clear evidence for selection in favor of a larger hypomethylated region (*dM* = 604.15). The average hypomethylated region width is 4403 for the top 10% loss-of-function intolerant gene promoters, in contrast to 2897 for the bottom 10%, with a gradual shift in betweeen (Figure 2b; *p* = 9.9 · 10^-5^).

### DNA methylation at the transcriptional end region in the male germline is not under selection

DNA methylation at proximal promoters has been studied extensively, because of its association with gene expression. Much less attention has been paid to methylation at transcriptional end regions, even though it has been recognized that the 3’ ends of some genes can also be hypomethylated (Gardiner-Garden, Frommer, 1987; Yu et al., 2013). The functionality of variation in transcriptional end region methylation remains unknown, prompting us to ask if selection is at play (Methods). We treated transcriptional end region methylation as a binary epigenetic mark (Supplemental Figure S4c; Methods), and did not find any evidence for selection (*dM* = 0.0004). The percentage of methylated transcriptional stop sites was essentially constant across LOEUF deciles, with minor fluctuations that did not deviate from what would be expected by random chance alone (Figure 2c; *p* =0.45), in concordance with what our test predicts under neutrality.

### The presence of H3K36me3 at coding regions in the male germline is under positive selection

We next shifted our focus to a histone modification: gene-body H3K36me3. Using ChIP-seq data from human sperm, we called peaks genome-wide and subsequently restricted to peaks that overlap coding regions (Methods). With those peaks, we treated H3K36me3 as a binary mark that is either present or absent in the coding region. We found that greater loss-of-function intolerance is associated with a higher probability of an H3K36me3 peak in the coding sequence (Figure 2d; *dM* =0.09; p = 9.9 · 10^-5^), showing that gene-body H3K36me3 is under positive selection in the male germline.

### The presence of H3K4me3 and the intensity of the H3K4me3 signal at proximal promoters in embryonic stem cells are under positive and directional selection, respectively

Finally, we examined proximal promoter H3K4me3, using ChIP-seq data from human H1 embryonic stem cells (Methods). We first tested for selection on the presence of at least one H3K4me3 peak at the proximal promoter. We found clear evidence for positive selection (Figure 2e; *dM* = 0.12; *p* = 9.9 · 10^-5^). This is not unexpected, as having an H3K4me3 peak at the promoter has been strongly linked to gene regulation. However, it is not clear whether, across promoters which harbor such a peak, differences in the intensity of the ChIP-seq signal (Supplemental Figure S4d) reflect underlying biology or technical noise (Landt et al., 2012; Marinov, Kundaje, 2018). Applying our test to H3K4me3 signal intensity can provide an unbiased answer to this question, as a result indicative of selection would imply that the intensity of this signal reflects biologically meaningful regulatory activity. Indeed, our test revealed the action of directional selection favoring a higher intensity, with a *dM* value of 4.48 (Figure 2f; *p* = 9.9 · 10^-5^). We emphasize that the signal intensity here is merely a read-out; selection acts on the biological process which produces this signal, likely the kinetics of H3K4me3 deposition/removal.

### Gene-specific optimization of DNA methylation and H3K4me3 at proximal promoters is unlikely

For many epigenetic marks, we observe different states throughout the genome. For example, some proximal promoters are methylated while others are hypomethylated. Under our model, where each *trans*-regulatory group has its own fitness and the mark is under the same kind of selection (positive or negative) at all groups, different states can still arise if selection is not strong enough to overcome drift at some groups. In this case, a different state than the one favored by selection may reach fixation via random drift. An alternative possibility, however, is that different states arise because the mark is under negative selection at some groups but under positive selection at other groups; in other words, there is location-specific “optimization” of the epigenetic state. We sought to investigate this possibility. From the epigenetic marks we tested above, we focused on the two that are arguably the most well-understood: the presence of DNA methylation and H3K4me3 at proximal promoters.

Given the known relationship between proximal promoter DNA methylation and gene repression (XS Liu et al., 2016; Korthauer, Irizarry, 2018), we reasoned that if DNA methylation at the proximal promoter has been favored by selection at some groups of genes, then these would be genes where a reduction in expression was beneficial. Therefore, these genes now should not tolerate an increase in their expression levels (assuming that the selective pressure on their expression level is conserved). To examine if this is the case, we first used recent estimates of “triplosensitivity” (Collins et al., 2021). These estimates capture the extent to which genes tolerate increased expression. We found that the genes whose proximal promoters have DNA methylation are not triplosensitive (Supplemental Figure S5; median probability of triplosensitivity = 0.19). In fact, the genes whose proximal promoters are devoid of DNA methylation are the ones with high triplosensitivity estimates (Supplemental Figure S5a); this is consistent with the positive correlation between triplosensitivity and *s*_het_ estimates (Collins et al., 2021). The same reasoning applies to the absence of H3K4me3 at proximal promoters, which is also associated with low expression. Genes whose proximal promoters are devoid of H3K4me3 are not triplosensitive (Supplemental Figure S5b; median probability of triplosensitivity = 0.17), whereas those that do harbor H3K4me3 are (as expected given the association between promoter hypomethylation and H3K4me3; Ooi et al. (2007))).

To corroborate these results with an orthogonal approach, we investigated expression quantitative trait loci (eQTLs) in early human development (induced pluripotent stem cells; DeBoever et al. (2017); Methods) and male germline (testis; GTEx Consortium (2020); Methods). We found that, out of the mapped eQTLs which increase expression levels, the eQTLs corresponding to genes with methylated proximal promoters confer higher increases (Supplemental Figure S5c, d). This is the opposite of what would be expected if these genes were intolerant to expression level elevation. For H3K4me3, there exists no difference between the increase in expression conferred by eQTLs of genes harboring H3K4me3 peaks at their proximal promoter, and the increase conferred by eQTLs of genes without H3K4me3 (Supplemental Figure S5e); this is again inconsistent with the existence of greater selective pressure against expression level elevation at genes without H3K4me3.

Taken together, these data suggest that the presence of DNA methylation and the absence of H3K4me3 at proximal promoters are unlikely to have resulted from positive selection.

### The selective pressure on the epigenetic marks tested is not purely a passive consequence of selection on gene expression

Since the early days of the epigenetics field, there has been intense debate about whether epigenetic marks actively dictate cellular functions, or whether their observed patterns along the genome are purely a passive byproduct of gene expression regulation by other means, such as transcription factors (Cooper, 1983; AP Bird, 1984; Ptashne, 2007; Murray et al., 2019; Z Wang et al., 2022). This is a specific instance of a more general problem: unlike the DNA sequence, intermediate molecular features such as epigenetic marks can be affected by other features, such as gene expression levels. Therefore, one has to consider whether the inferred selective pressure on a certain feature is an artifact of selection on another feature. Our framework allows us to evaluate this for *a priori* chosen features, by recognizing and leveraging the connection between our setup for selection inference and causal mediation analysis (Figure 3a; Methods) (Pearl, 2014; VanderWeele, 2016).

**Figure 3.**
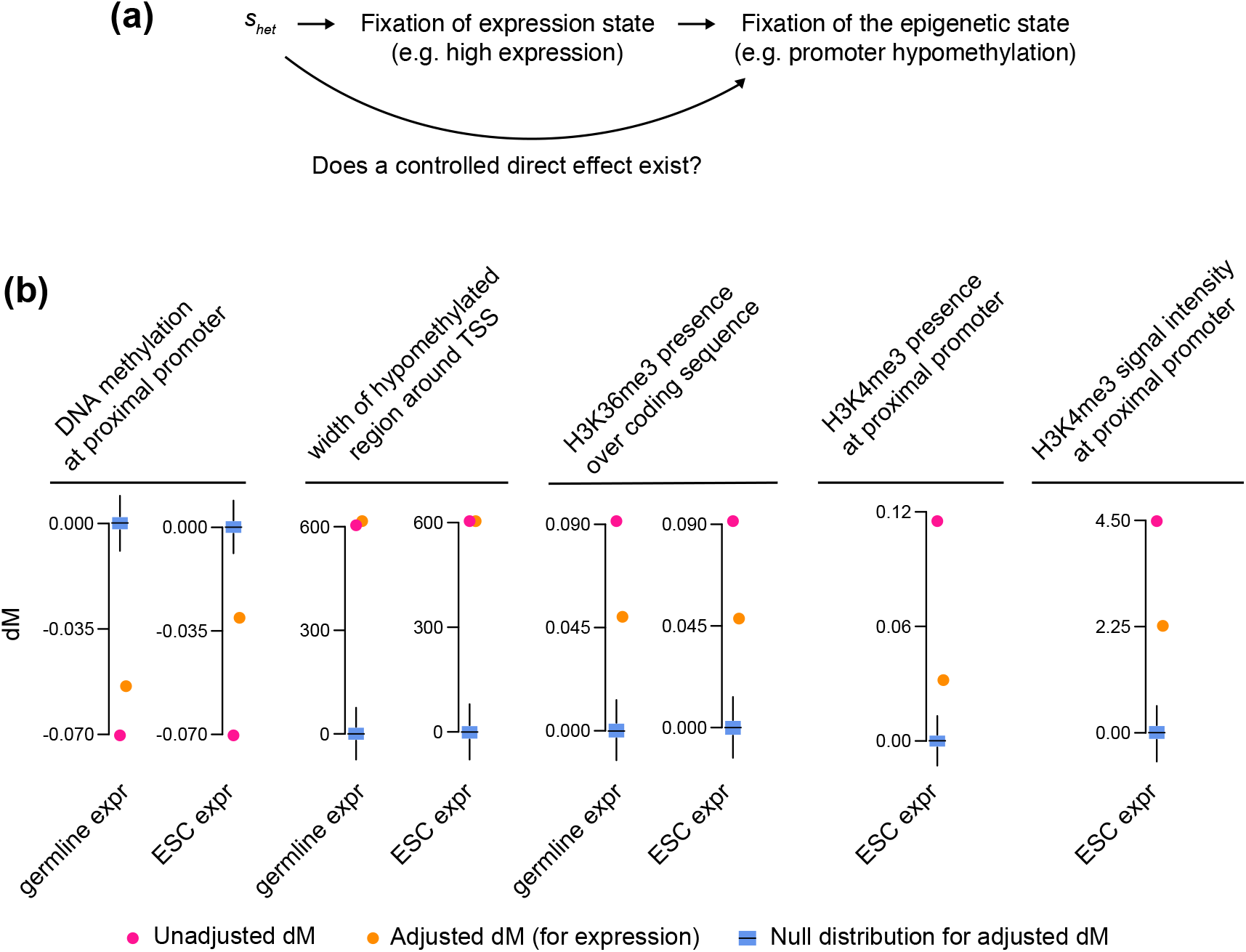
The selective pressure on the epigenetic marks is not fully explained as a passive consequence of selection on gene expression. **(a)** Hypothetical causal graph depicting the relationship between *s*_het_, the fixation of a specific gene expression state, and the fixation of the epigenetic state. We wish to test if the fixation of the epigenetic state is purely a passive consequence of selection on gene expression, i.e. whether or not there exists a controlled direct effect from *s*_het_ to the fixation of the epigenetic state. **(b)** For different marks, the observed values for the *dM* test statistic (red dots) and the corresponding null distributions (blue boxes), after adjusting for gene expression in the germline or human H1 embryonic stem cells. The unadjusted *dM* values from Figure 2 are included for comparison.

We thus set out to test if the selection signal on the epigenetic marks we investigated is a passive consequence of selection on gene expression. We discovered that this is not true. First, for the germline marks, we computed the adjusted dM after taking the effect of germline gene expression into account. In all cases, the observed adjusted *dM* values remain statistically significant (Figure 3b, d, f). Following mediation analysis (Methods), this is evidence for a direct effect of the mark on fitness, provided the noise in expression measurements is not too high. We then investigated the possibility that these germline epigenetic patterns are established as a consequence of expression patterns not in germ cells, but in subsequent developmental stages post-fertilization, and are merely inherited unaltered by primordial and subsequently mature germ cells. As representative of early pre-implantation development preceding primordial germ cell specification, we examined embryonic stem cells (bulk RNA-seq; Methods). For completeness, we also looked at subsequent fetal development (172 fetal cell types identified using single-cell RNA-seq; Methods), in case some of these transcriptional profiles correlate better with the primordial germ cell profile. In all cases, the adjusted *dM* values were still consistent with selection (Figure 3b; Supplemental Figure S6). The same is true for H3K4me3 presence and intensity, after adjusting for expression either in embryonic stem cells, or in fetal cell types (Figure 3b; Supplemental Figure S6).

### However, a causal role of epigenetic marks in gene expression regulation may not entirely explain their selective pressure

The results in the previous section are incompatible with the notion that the selection signal on the epigenetic marks is entirely explained as a passive consequence of selection on gene expression. A straightforward alternative explanation is that these marks contribute actively to normal cellular physiology and ultimately organismal fitness, by exerting a causal influence on gene expression regulation. Such a causal influence is supported by experimental evidence, primarily from targeted epigenome editing (XS Liu et al., 2016; Bintu et al., 2016; Korthauer, Irizarry, 2018; XS Liu et al., 2018; Mendoza et al., 2022). However, this interpretation raises the question: is expression regulation the only reason for selection on epigenetic marks?

To investigate this, we focused on germline promoter DNA methylation and its relationship with expression. First, we observed that the correlation between gene expression and the width of the hypomethylated region around hypomethylated proximal promoters is extremely weak, both in the germline and in embryonic stem cells. Hypomethylated region width in the germline is not significantly associated with germline gene expression (*p* = 0.7), whereas it only accounts for a negligible 0.1% of the variance in expression in embryonic stem cells (Supplemental Figure S7a, b). This suggests that expression regulation is unlikely to be the reason that selection acts on the the width of the hypomethylated region. Second, we identified 36 genes whose proximal promoters are methylated in the germline and become hypomethylated in embryonic stem cells (Methods). As a group, these genes are less intolerant to loss-of-function variation than genes with proximal promoters hypomethylated in both germline and embryonic stem cells (Supplemental Figure S8a), but they are expressed at similar levels (Supplemental Figure S8b). Although this is a very small set of genes, their existence illustrates that – even at proximal promoters – germline hypomethylation is preferentially targeted away from loss-of-function intolerant genes in the germline, without this being an absolute requirement for high expression in early embryonic stages. This is consistent with recent results from Mendoza et al. (2022), showing that forcible promoter hypermethylation is not necessarily accompanied by gene repression; there are genes whose expression is unchanged, and even some whose expression increases.

Finally, a general argument applicable to all epigenetic marks is that they do not exercise their effect on gene regulation alone. Rather, that effect is mediated by other components of the cellular epigenetic machinery, such as methyl-CpG binding proteins and histone modification readers (Jørgensen, A Bird, 2002; Musselman et al., 2012; Boukas et al., 2019). It is unlikely that this multi-component system evolved at once. One plausible scenario is that the marks appeared first, were retained by selection unrelated to gene expression (as well as by drift), and then the protein components appeared, creating a functional gene regulatory system.

### Can the effect of DNA methylation and H3K36me3 on regional mutation rates partly explain their selective pressure?

Motivated by the above observations, we searched for explanations of selection on epigenetic marks unrelated to expression. An intriguing hypothesis pertaining to DNA methylation and exonic H3K36me3 is that these marks are under selection partly because they affect regional mutation rates. We set out to test the basic premises of this hypothesis, and investigate if such secondary selection can overcome random genetic drift in a finite population.

The basic premises of the hypothesis are that: a) these marks indeed cause changes in mutation rates, and b) the resulting mutations are deleterious. It is well-established that DNA methylation increases the mutation rate of CpGs in the germline by ~3-4 fold (Coulondre et al., 1978; RY Wang et al., 1982; Cooper, Youssoufian, 1988; Zhou et al., 2020). However, while we have previously observed a strong link between CpG density at promoters and genic intolerance to loss-of-function mutations (Boukas et al., 2020a), it is unclear if the loss of CpGs at promoters has fitness consequences, as previous studies have reached conflicting conclusions (Cohen et al., 2011; Panchin et al., 2016). For exonic H3K36me3, on the other hand, it is well-accepted that the resulting mutations are deleterious on average, since they are coding. But, while H3K36me3 has been found to promote mismatch repair and homologous recombination repair in somatic cells (Li et al., 2013; Pfister et al., 2014; Z Sun et al., 2020), it is unknown whether its presence is associated with a reduced exonic mutation rate in the germline. A previous study found no such association, in contrast to the situation in somatic cells (Rodriguez-Galindo et al., 2020). However, this study used H3K36me3 from embryonic stem cells as a proxy for the germline profile, and therefore did not resolve the question.

### Higher loss-of-function intolerance is associated with lower *de novo* coding mutation rates, an effect partly explained by H3K36me3

Given that higher genic loss-of-function intolerance is associated with a higher probability of exonic H3K36me3, we started by asking whether higher loss-of-function intolerance is also linked to a lower exonic mutation rate. We used a dataset of 15,642 *de novo* synonymous mutations, aggregated from 58,011 individuals (Methods; Zhao et al. (2020)). This revealed that, indeed, the more loss-of-function intolerant a gene is, the lower its exonic mutation rate (Figure 4a; Supplemental Figure S9a). Genes within the top 10% of loss-of-function intolerance have on average 0.0007 de novo synonymous mutations per coding sequence base pair. By contrast, the corresponding average rate for genes within the bottom 10% of loss-of-function intolerance is 0.0012, that is, 1.71 times higher. We note that the use of *de novo* synonymous mutations ensures that this result truly reflects differences in mutation rates, and not the action of selection. We found that including intronic mutations makes the association between loss-of-function intolerance and textitde novo mutation rate a lot weaker (Supplemental Figure S9e,g,h), emphasizing that this phenomenon is largely localized to the regions under selection: exons.

**Figure 4.**
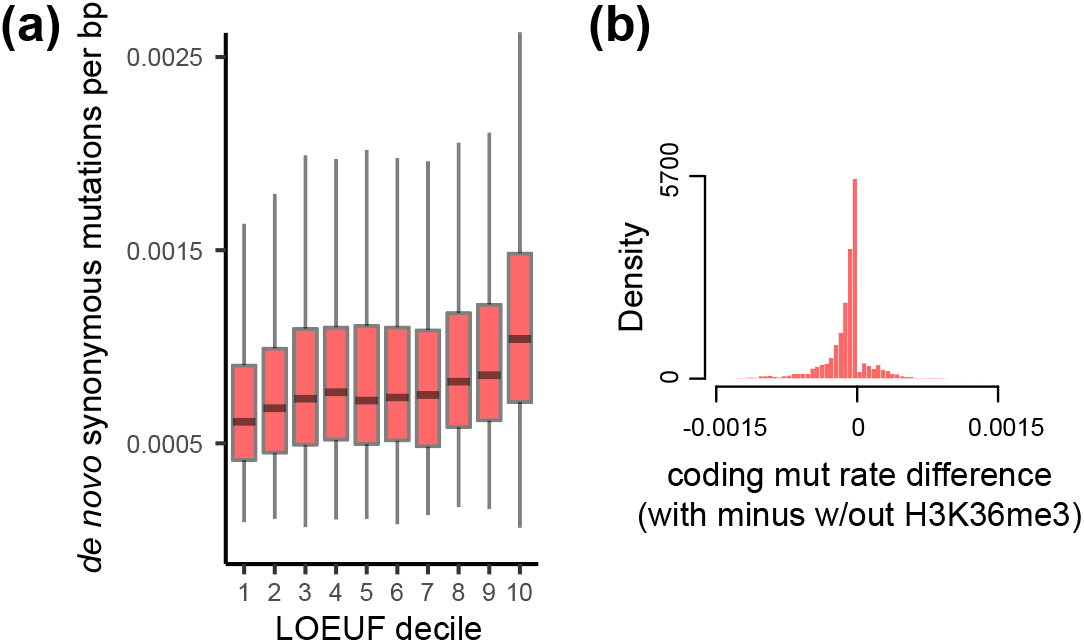
Higher loss-of-function intolerance is linked to lower *de novo* coding mutation rate, partly explained by coding H3K36me3 patterns. **(a)** The distribution of the number of synonymous *de novo* mutations per coding sequence base pair, stratified based on downstream gene loss-of-function intolerance. **(b)** The distribution of the difference between the number of synonymous *de novo* mutations per coding sequence base pair in H3K36me3-marked regions and regions without H3K36me3. H3K36me3-marked regions are defined as a 500bp interval centered on each H3K36me3 peak (restricting to exonic regions only). The distribution is obtained by computing the difference for each gene.

We next assessed the contribution of H3K36me3 to this pattern. Clearly, since only 40.3% of genes have an H3K36me3 coding peak, H3K36me3 cannot be the sole determinant of the association between exonic mutation rate and intolerance to loss-of-function mutations. Nevertheless, we focused on genes with at least one H3K36me3 coding peak and recalculated the number of *de novo* synonymous mutations per base pair, this time after excluding the sequences directly within or very close (+/− 250bp) to H3K36me3 peaks. Comparing this rate to the corresponding one when using the entire exonic sequence, we found that 86.2% of genes have a higher mutation rate after excluding H3K36me3-associated sequences (Figure 4b; Supplemental Figure S9b). This confirms that germline H3K36me3 is associated with lower exonic mutation rates, and shows that this association is present in the vast majority of genes which harbor an H3K36me3 coding peak.

### Promoter CpGs at loss-of-function intolerant genes are under selection

Turning our attention to CpGs, we investigated if their loss at promoters is deleterious or not. We used conservation across 100 vertebrate species, quantified using the PhyloP score (with higher values indicating greater conservation; Methods). We examined 1,076,680 proximal promoter CpGs and 563,098 CpGs that are in hypomethylated regions but not within proximal promoters (which we refer to as promoter boundary CpGs).

While conservation alone generally does not necessarily reflect selection, conservation at promoter nucleotides does reflect selection if it increases when genic loss-of-function intolerance increases. We found that the per promoter average CpG conservation increases as downstream gene loss-of-function-intolerance increases. This is true both at the proximal promoter and at promoter boundaries (Figure 5a, c). Furthermore, this increase does not reflect background selection due to linkage to other promoter or coding sites, because it is accompanied by a progressively larger difference between the average per promoter conservation of CpGs versus that of non-CpG sites (Figure 5b, d).

**Figure 5.**
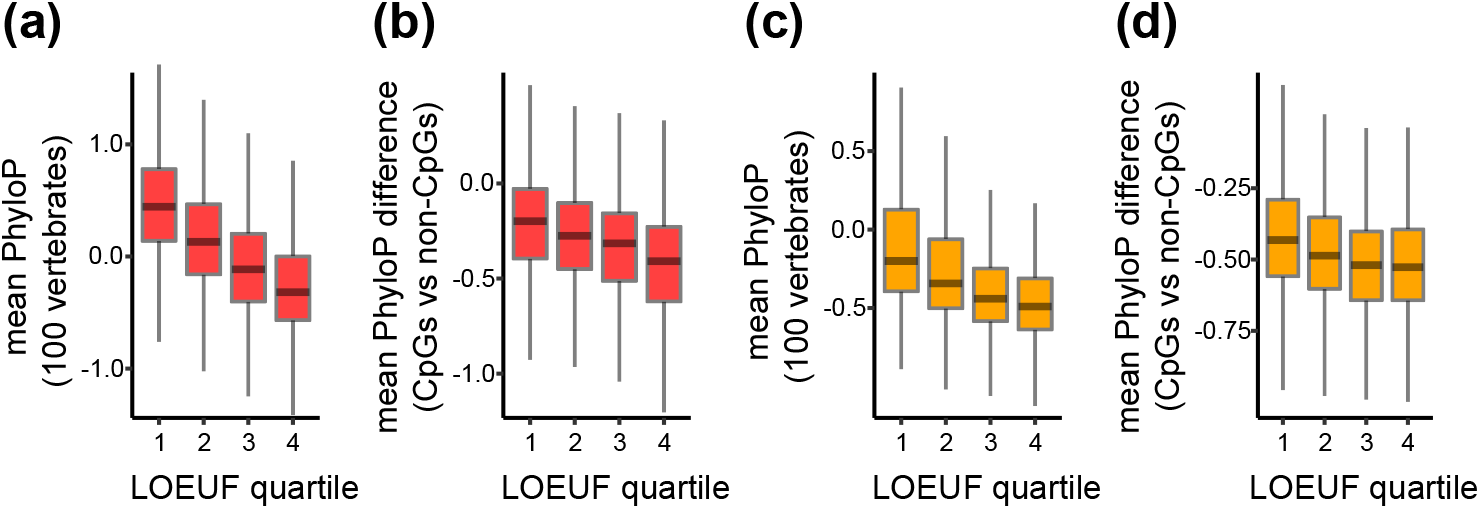
Higher genic loss-of-function intolerance is associated with increased conservation of hypomethylated promoter CpGs. **(a)** The distribution of per promoter average across-vertebrate conservation (PhyloP score) of CpG sites in the proximal promoter, stratified according to downstream genic loss-of-function intolerance. **(b)** The distribution of per-promoter differences in the average conservation (PhyloP score) of CpG sites minus that of non-CpG sites, stratified according to downstream genic loss-of-function intolerance. **(c,d)** Like (a,b) respectively, but for CpGs in the promoter boundary.

### Theory and simulations support the existence of selection on promoter CpG methylation and exonic H3K36me3 because of their effect on regional mutation rates

The drift-barrier hypothesis posits that – under realistic mutation rate values – the fitness effect of local changes in mutation rate is not strong enough to overcome random drift (Lynch, 2011; Lynch et al., 2016) in a finite population. However, the major feature of epigenetic marks, which has been central to our study, is their regulation in *trans* at multiple genomic loci. This implies that changes in epigenetic state will affect multiple locations simultaneously. In light of this, we re-evaluated the drift-barrier hypothesis. We based our investigation on a well-established model for the accumulation of deleterious mutations at genomic regions (Methods; Haigh (1978)). While this model does not capture the full complexities of biological reality (see Methods for simplifying assumptions used), we reasoned that it can provide useful insights.

To make our results as realistic as possible, we used empirical estimates for selection coefficients of mutations in conserved non-coding and coding regions (Kryukov et al., 2005). In addition, we empirically estimated the relevant mutation rates (Methods) and, in the case of DNA methylation, the proximal promoter “epimutation” rates (via a human-chimp comparative analysis of sperm methylomes with rhesus as an outgroup; Methods). Of particular note, we found these “epimutation” rates to be 3.23 · 10^-8^ for methylated promoters and 2.4 · 10^-9^ for hypomethylated ones. These values are on par with and lower than, respectively, mutation rates in human coding sequences. Therefore, because we are considering marks occurring at multiple locations throughout the genome, we hereafter neglect this epimutation rate, in analogy with the assumption underlying the standard 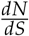 test for coding sequences (Nielsen, Yang, 2003; Kryazhimskiy, Plotkin, 2008). An analysis of CpG genetic diversity among 62,874 individuals supports this approximation (Supplemental Figure S11; Methods, see also last paragraph of section “A test statistic for selection”). This immediately implies, that in an infinite population, the epigenetic state associated with lower mutation rate will eventually dominate in the population (Methods).

Turning to the case of a finite population, we first examined the limiting case of a small region in a single gene (corresponding, for example, to a single promoter or a single H3K36me3 exonic peak). In agreement with the drift-barrier hypothesis, we observe that the difference in mutation rate it too small for selection to overcome drift (Figure 6); the fixation probability is the same as what is expected under neutrality. We corroborated this simulation result with a heuristic argument based on a diffusion approximation to the probability of the epigenetic mutation rate modifier (Methods).

**Figure 6.**
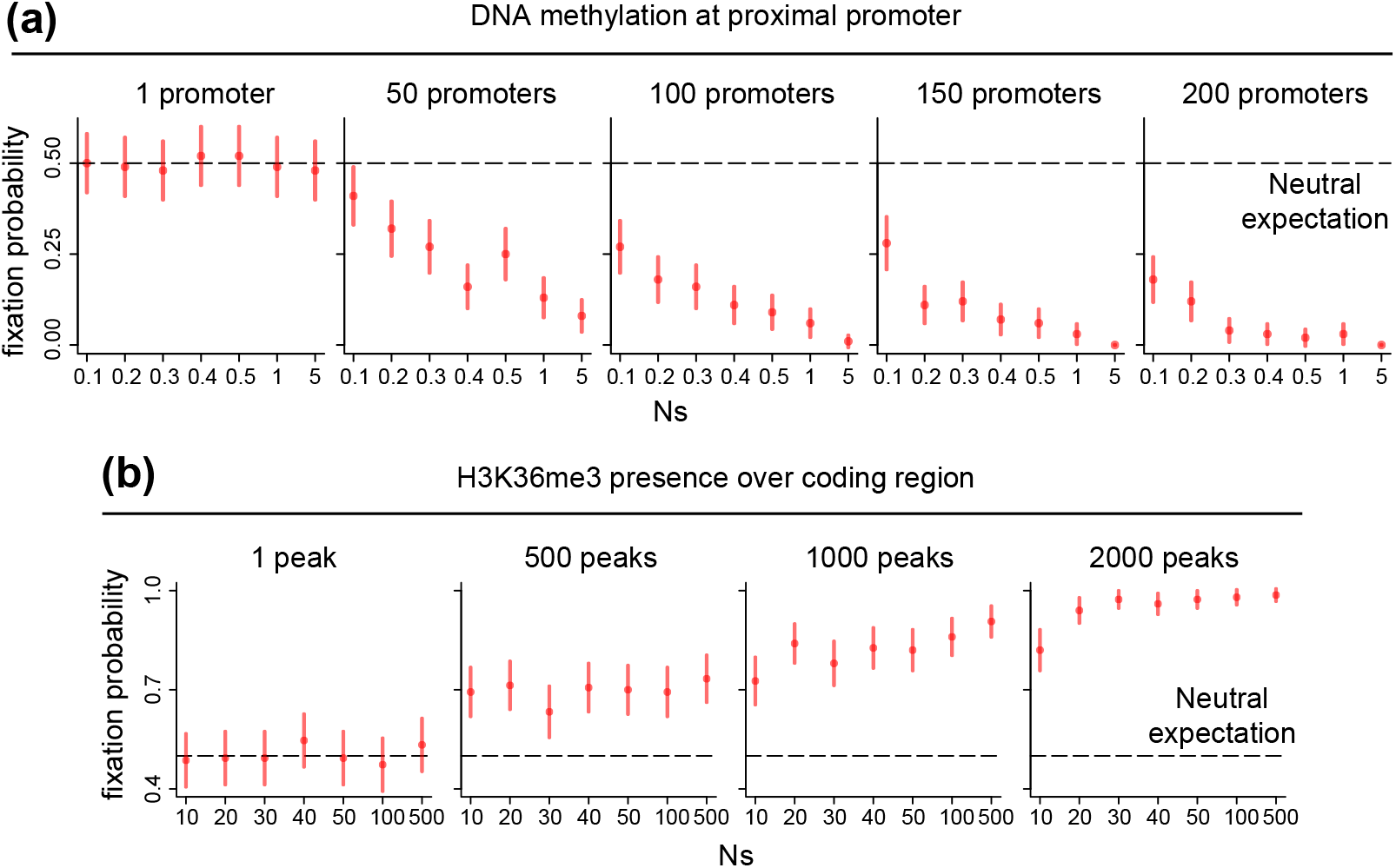
The selective pressure on promoter DNA methylation and coding H3K36me3 may be due to their effect on regional germline mutation rates. **(a)** The probability of fixation of proximal promoter DNA methylation, under different scenarios. Each scenario corresponds to a different value of the product between the effective population size and the selection coefficient (*Ns*) and to different numbers of promoters under simultaneous control by upstream *trans*-regulators, with 80 CpGs per promoter (Methods). **(b)**. The probability of fixation of H3K36me3 at the coding region. As in **(a)**, different scenarios are depicted, corresponding to different values of Ns and the numbers of H3K36me3 peaks under simultaneous control.

We then examined different scenarios with increasing number of regions under upstream regulatory control. We observed that, as the number of regions under upstream control increases, the effectiveness of selection increases as well (Figure 6). For DNA methylation, the increase in CpG mutation rate is such that even with small *trans*-regulatory groups, selection overcomes drift (Figure 6). Moreover, the fixation probability decreases when the selection coefficients increase (Figure 6), a trend which qualitatively mirrors our empirical results. For exonic H3K36me3, even though the effect on the mutation rate is smaller, selection still becomes effective when the *trans*-regulatory group size increases, although the required number of regions is substantially higher than for DNA methylation (Figure 6). This is not necessarily inconsistent with biological reality, as coding H3K6me3 patterns are largely set by the globally-acting SETD2 histone methyltransferase (XJ Sun et al., 2005; Bhattacharya, Workman, 2020). Again, we observe a gradient of decreasing fixation probabilities with increasing selection coefficients (Figure 6), although this gradient is less pronounced than in the case of DNA methylation.

These results show that *trans*-regulation of multiple regions can make regional epigenetic mutation rate modifiers subject to selection strong enough to overcome drift. This can reconcile our observations with population genetics theory.

## Discussion

We have presented a framework for detecting natural selection on epigenetic marks. Our main contribution is twofold. First, we show that estimates of selective pressure against heterozygous coding loss-of-function alleles – which have recently become available – motivate a definition of the fitness effect of epigenetic marks associated with genes. Second, we use this definition to derive a test for selection. Ultimately, we provide a characterization of what it means for an epigenetic mark to be under selection versus not. If a mark is equally likely to be encountered across genes regardless of their tolerance to loss-of-function variation, it is not under selection; if it less (more) likely to be encountered the more loss-of-function intolerant a gene is, the mark is under negative (positive) selection.

More fundamentally, our framework is general. There is nothing that restricts its applicability to epigenetic marks only, and we envision that several molecular (and genomic) features will be amenable to testing with the same approach. This rests on the belief that the central function of the genome is to transcribe genes. Thus, a test for selection which relies on a genic *s*_het_-based fitness definition will be appropriate for a multitude of features. Importantly, our framework incorporates adjustment for confounders, an essential component of any test for selection on intermediate molecular features. We use this to establish that selection on the marks we consider is not merely a passive consequence of selection on gene expression. With any given feature, however, one should carefully decide what confounders to adjust for, and this choice must be informed by the relevant biology. This illustrates a caveat of our approach, as it may often be difficult to completely rule out the possibility that an inferred selective pressure is driven by confounders that were not accounted for. Another limitation behind our test currently is that it provides information about the presence or absence of selection, but not about its strength. Future work should address this gap, although it may require the full specification of how a mark is regulated in *trans*.

We note that the core idea underlying our test and its application here - the assessment of the relationship between genic loss-of-function intolerance and the presence of an epigenetic mark throughout the genome - may seem intuitively obvious. Indeed, an initial version of our work, focusing only on germline promoter DNA methylation (Boukas et al., 2020b), as well as work by (Monroe et al., 2022) on A. Thaliana, use this idea in a heuristic fashion. Here we make the approach precise, which illustrates its generality and the subtleties underlying its application. In addition, we provide a framework based on the notion that epigenetic marks are controlled by *trans*-acting regulators. This is a major conceptual issue that was ignored in previous studies, and - as we also discuss below - takes center stage when assessing the impact of epigenetic marks on mutation rates, the subject of the aforementioned studies.

Our results also yield biological insights. First, with regards to specific marks, probably what is most unexpected is the positive selection detected on the width of the hypomethylated region around hypomethylated proximal promoters. This is despite the fact that there is a negligible association between the size of this region and gene expression. These hypomethylated regions largely correspond to CpG island shores (Irizarry et al., 2009), which are characterized by rapid methylation-demethylation kinetics (Ginno et al., 2020), and are often differentially methylated between tissues. Whether these properties are relevant to the selective pressure we detect here in the germline remains to be determined.

Second, more generally, whether the epigenetic states encountered in the vicinity of genes are purely a passive byproduct of gene expression regulation with no active role of their own has long been debated (Cooper,1983; AP Bird, 1984; Ptashne, 2007; Murray et al., 2019; Z Wang et al., 2022). To the best of our knowledge, our study is the first to approach this question through an evolutionary lens. Our findings show that epigenetic marks have a causal effect on cellular function strong enough to render them subject to selection.

It is tempting to conclude that the epigenetic marks considered here are under selection because of their involvement in gene regulation. Indeed, targeted epigenome editing experiments support this interpretation (XS Liu et al., 2016; Bintu et al., 2016; Korthauer, Irizarry, 2018; XS Liu et al., 2018). However, here we make the case that aside from a causal role in expression regulation, complementary explanations are also needed in order to explain the full selective pressure. This is motivated in part by the fact that some aspects of promoter DNA methylation are only weakly correlated with expression, and in part by a general consideration concerning the interplay between epigenetic marks and their readers.

For promoter DNA methylation and exonic H3K36me3, a complementary explanation could be their impact on local mutation rates in the germline. Whether selection acts on local mutation rate modifiers is a very old question in population genetics and molecular evolution. A long line of theoretical investigations has led to the proposal that selection may act to reduce the mutation burden at location where mutations are deleterious (Sturtevant, 1937; Kimura, 1967; Leigh, 1970; Kondrashov, 1995). However, empirical evidence for this phenomenon has been lacking. In fact, it has been questioned whether such secondary selection can overcome the force of random genetic drift in finite populations (Lynch, 2011; Lynch et al., 2016), an argument termed the drift-barrier hypothesis. Our simulations here show that – under realistic mutation rates and selection coefficients − the drift barrier can be overcome in the case of epigenetic mutation rate modifiers that are under trans-regulation, because they affect multiple genomic regions simultaneously. This provides support for the notion that regional “optimization” of promoter and coding germline mutation rates has shaped the evolution of DNA methylation and H3K36me3, and is relevant for the interpretation of a recent result in *A. thaliana* (Monroe et al., 2022) as well. But we highlight that it is is non-trivial to answer conclusively whether there is selection on a mark because of its effect on local mutation rates, on top of selection due to a causal effect on gene expression that the mark has at the same time. At best, one can establish – as we do here – that the secondary selection due to mutation rate modification is consistent with evolutionary theory. Providing “proof” for this is likely to require experimental evidence with directed evolution (Morselli et al., 2015; Finnegan et al., 2020). However, even for DNA methylation, whose effect on the mutation rate is strong and unequivocally established, it will be difficult to retain that effect while simultaneously blocking its potential effect on gene expression, since the mere methylation of CpGs changes the local thermodynamic properties of DNA (S Wang, Kool, 1995; Renciuk et al., 2013; Tsuruta et al., 2021) and can modulate transcription factor binding (Yin et al., 2017; Héberlé, Bardet, 2019).

It is worth commenting further on the potential for selective pressure on CpGs via the exclusion of DNA methylation. Earlier work on the evolution of CpG islands in primates reported no evidence for selection maintaining them (Cohen et al., 2011). This work has become accepted by leading members of the epigenetics community (Antequera, A Bird, 2018). However, the approach in Cohen et al. (2011) assumes that, when studying the evolutionary dynamics of promoter CpGs, DNA methylation is merely a confounder that needs to be adjusted for. This rests on the long-standing, widely held belief that promoter CpG-richness is a passive consequence of the hypomethylated state of these promoters in the germline. Our work provides evidence for the reverse rationale: that part of the reason why these promoters have remained hypomethylated is because this is how selection is preserving their CpG-richness. Such a scenario is consistent with the high mutational constraint of CXXC-domain-containing proteins, which bind to unmethylated CpGs (Lee et al., 2001; Boukas et al., 2019) and are also present in organisms lacking DNA methylation FIXME: cite, and with the role of these unmethylated CpGs in expression regulation (Thomson et al., 2010; Clouaire et al., 2012; Hartl et al., 2019).

Finally, we have shown that the more intolerant to loss-of-function variation a gene is, the lower its exonic mutation rate in the germline. In light of a recent similar result in *A. thaliana* (Monroe et al., 2022), and previous observations in *E. coli* (Martincorena et al., 2012), it is natural to suggest that this is a universal phenomenon: essential genes have an inherently lower propensity to mutate across diverse organisms. We emphasize that while we here focus on the contribution of H3K36me3, H3K36me3 is certainly not the sole factor explaining the variation in has the potential to provide new clues into the determinants of germline mutation rates. Of relevance here may be the recently proposed “transcriptional scanning” hypothesis (Xia et al., 2020; Xia, Yanai, 2022).

## Methods

### A model for the fitness effect of an epigenetic mark regulated in *trans*

We consider an epigenetic mark which occurs at multiple sites throughout the genome. We restrict our attention to cases where each individual site is associated with a specific gene. Examples of such sites where epigenetic marks are known to occur are promoters and gene bodies. Hereafter, we use the words site and gene interchangeably.

For simplicity, we assume that at each gene there are two possible epigenetic states, which we call *M* and *U*. As we show later, our test for selection is readily applicable to marks whose states vary along a continuous spectrum.

### *trans*-regulatory groups

As described in the Results section, epigenetic marks are regulated in *trans* by upstream factors. Owing to this, the same epigenetic state is present at a group of genes which are under shared upstream regulation. What we observe at an individual gene is a consequence of the particular *trans*-regulatory group that the gene belongs to. To capture this *trans*-regulation in our model, we denote a given trans-regulatory group by *G*, and use *w_G_* to denote the fitness of individuals with the *M* state at this group. This fitness *w_G_* will be a function of the fitness effect of the *M* state at each of the genes belonging to group *G*.

Before providing a model for this fitness effect at the single-gene level, let us further illustrate the reasoning behind a *trans*-regulatory group, with a simple scenario. Consider a single transcription factor that binds to a set of promoters and recruits chromatin modifiers, which in turn set up a particular epigenetic state. We assume that, when expressed and active, the transcription factor binds to all promoters that contain its cognate motif. In that case, a group *G* consists of the set of genes whose promoters harbor the transcription factor motif. At a given individual and within a given cell type, the epigenetic state we observe at these genes is determined by whether the transcription factor is expressed and is active, or not. Variation across the population in that cell type is governed by variation in the expression and activity of the transcription factor. However, the networks of upstream regulators (transcription factors and chromatin modifiers) and their interactions, which define these groups, are currently incompletely characterized.

#### Example (DNA methylation)

We now briefly describe a concrete example relevant to the regulation of DNA methylation provided in Boulard et al. (2015). In that study, the focus is on KDM2B, a regulatory factor (histone demethylase that also has DNA and histone binding ability) which binds to approximately 15,000 CpG-rich promoters via its CXXC domain. It was shown that knockout of KDM2B in embryonic stem cells leads to a gain of DNA methylation, but only at a subset of CpG-rich promoters that are also bound by Polycomb repressive complexes 1 or 2. This implies the existence of the following regulatory groups: a) the group of genes whose promoters are bound by both KDM2B and Polycomb repressive complex 1 or 2; these promoters are unmethylated when KDM2B is expressed at normal levels; b) the group of genes whose promoters are not bound by KDM2B; these promoters are normally methylated; c) the group of genes whose promoters are bound by KDM2B, but not by Polycomb repressive complex 1 or 2; these genes are in fact probably split into more than one regulatory group, and the logic determining their DNA methylation state is unknown.

### From the fitness effect on a single gene to the fitness effect on a trans-regulatory group

We now present an explicit model that connects the fitness effect of a given epigenetic state at individual genes with the resulting aggregate effect on a *trans*-regulatory group. In doing so, we make explicit the contribution of genic *s*_het_ coefficients, which is critical for our subsequent inference procedure.

#### A single gene

At a given gene *g*, we define the fitness of an individual heterozygous for the *M* epigenetic state as

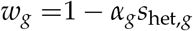

where *s*_het,*g*_ is the gene-specific selection coefficient against heterozygous coding loss-of-function alleles, and *α_g_* is a gene-specific parameter that is positive if *M* is under negative selection and negative if *M* is under positive selection (strictly speaking, the fitness should be defined as *w_g_* = max(0,1 – *α_g_s*_het,*g*_)).

*α_g_* can be thought of as a parameter that reflects how sensitive the gene is to alterations of its epigenetic state. For instance, in the case of negative selection, one may impose the additional restriction that 0 ≤ *α_g_* ≤ 1. Then, the closer *α_g_* is to 1, the more important the epigenetic mark is for the gene, and changes in the epigenetic state have almost as large of a fitness effect as a coding loss-of-function mutation (which provides an upper bound since it completely inactivates the gene by leading to non-sense mediated decay). By contrast, if is close to 0, then the gene is unaffected by changes in epigenetic state.

The key implication of this fitness definition is that, at a given gene, if an intervention were to be applied that would change *s*_het,*g*_ (e.g. manipulation of the environment that makes a gene very intolerant to loss-of-function mutations), then the fitness effect of the epigenetic state would also change. The larger *α_g_* is, the larger that change would be.

Finally, there is nothing specific about M being an epigenetic state here; the same definition could be used to capture the fitness effect of any other type of feature.

#### Multiple genes

Our inference procedure does not make any assumptions about the form of the function that relates the fitness effects of the *M* state at each of the individual genes of the group to the group-level fitness *w_G_*. However, it is illustrative to briefly consider the case of a multiplicative fitness landscape. Then, we have

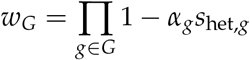

The set of (*α_g_*; *g* ∈ *G*) parameterizes the fitness of the *M* epigenetic state in terms of the set of (*s*_het,*g*_; *g* ∈ *G*). Importantly, these parameters only influence the evolutionary dynamics of the M state through their effect on *w_G_*.

### The fixation probability of an epigenetic state at a trans-regulatory group

An epigenetic state *M* at a *trans*-regulatory group *G* has a fixation probability which depends on the fitness *w_G_* (in addition to population size and starting frequency of the state). To emphasize this, we introduce the function *m* defined as

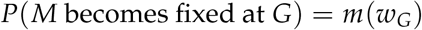

From standard population genetics theory, we know that, if the fitness effect is not strong enough, selection is dominated by random genetic drift in finite populations. We define *w_weak_* as a threshold such that

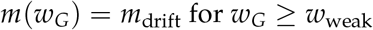

under negative selection, and

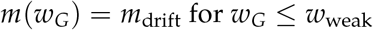

under positive selection.

This merely states that if selection cannot overcome drift, then the fixation probability does not depend on fitness (since it is governed by random fluctuations over time).

### A genome-wide model

We can divide the genome into disjoint (by definition) trans-regulatory groups *G*_1_,…, *G_k_* with associated fitnesses *w*_1_,…, *w_K_*. We assume without loss of generality that the groups are ordered in terms of these fitnesses, so that under neutrality or negative selection:

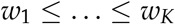

and under neutrality or positive selection:

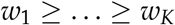

At each group separately, selection may (or may not) overcome drift, implying that the epigenetic state may be under selection only at some groups. We will say the epigenetic state is under selection if it is under selection for at least one *trans*-regulatory group.

### A quantity determined by selection

To construct a test for selection we need a quantity which behaves differently, i.e. attains different values, depending on whether the epigenetic mark of interest is under selection or not. Because the compoosition and logic of *trans*-regulatory groups are largely unknown, we need a quantity which does not depend on the composition of these groups. To obtain such a quantity, we examine the relationship between the probability of fixation and the gene-specific selection coefficients against heterozygous coding loss-of-function alleles *s*_het,*g*_, which are now known for most human genes (Cassa et al., 2017; Karczewski et al., 2020).

We consider the genome-wide model, where we have a finite set of genes with associated *α_g_* and *s*_het,*g*_. The genes are divided into *K* regulatory groups *G*_1_,…, *G_K_*. The presence of the *M* epigenetic state at these groups has an associated fitness *w*_1_ ≤ … ≤ *w_K_*.

We now consider the following experiment: we draw two genes independently at random (i.e. uniformly across genes) and start the evolutionary process. We define the following stochastic variables:

*g*, 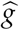 : The two genes
*S*, 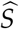 : The *s*_het_ coefficients of *g*, 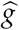 respectively
*M*^fix^, 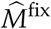 : Binary variables equal to 1 if the *M* epigenetic state is fixed at *g*, 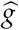 respectively

We define the following quantity:

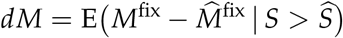

Our main result is the following, stated below for the case of negative selection (the case of positive selection is completely analogous and is treated afterwards):

#### Proposition 1.

*If the M epigenetic state is neutral or under selection that is too weak to overcome drift, i.e. if w_i_* ≥ *w_weak_, for all i, then*

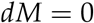

*If the M epigenetic state is under negative selection that is strong enough to overcome drift, i.e. there is at least one i such that w_i_* < *w_weak_ (and we do not have w*_1_ = ⋯ = *w_K_), then*

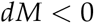

*under the assumption that differences in fitness between groups correspond to differences in s_het_ distributions, in the following sense:*

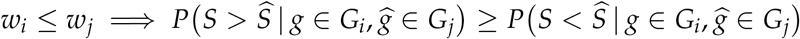

*with the inequality for the probabilities being strict when the inequality for the fitnesses is strict*

**Remark 1** The assumption describes first order stochastic dominance between certain conditional distributions. This is illustrated in Supplemental Figure S2 which considers 3 *trans*-regulatory groups where *w*_1_ > *w*_2_ > *w*_2_ together with *s*_het_, distributions under two different scenarios. It exemplifies that it is *possible* to have different fitness even with the same *s*_het,*g*_ distributions. As described in the “From the fitness effect on a single gene to a *trans*-regulatory group”, this different fitness effect can be caused by differences in the *α_g_* distributions.

**Remark 2** We emphasize that the assumption is only used in the second part of the theorem. When the epigenetic mark is neutral or under negative selection too weak to overcome drift, we always have *dM* = 0. When negative selection is strong enough to overcome drift, we have *dM* < 0, provided the assumption is satisfied. This means that observing *dM* < 0 is strong evidence for negative selection, whereas *dM* = 0 could follow either from lack of selection **or** violation of the assumption.

*Proof*. First, we rewrite

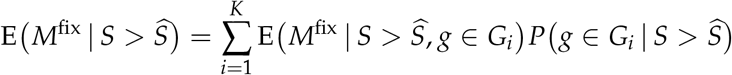

Conditional on {*g* ∈ *G_i_*}, and since we have a binary state, the expectation becomes the fixation probability *m*(*w_i_*) as previously discussed. So

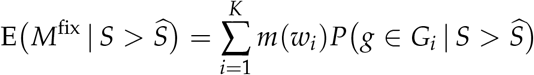

The same argument can be used to conclude

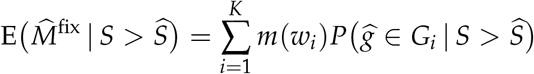

If *M* is neutral or selection is too weak to overcome drift, then *m*(*w_i_*) does not depend on *i*, and we can write *m*(*w_i_*) = *m*_drift_, for all *i*. We therefore conclude

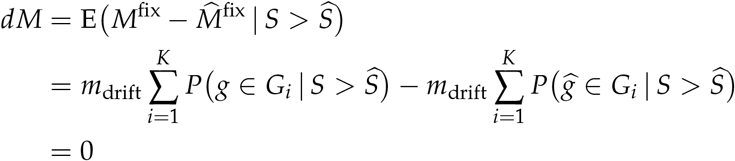

We now consider the negative selection regime, assuming that we do not have *w*_1_ = ⋯ = *w_k_*. Recall that without loss of generality, we assume that the groups are ordered according to the fitness effect of *M*. We therefore have that *w*_1_ < *w*_weak_. We return to our quantity of interest, *dM*:

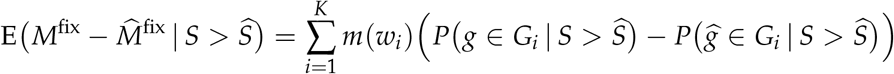

Now we use the identities (which are consequences of every gene belonging to a group)

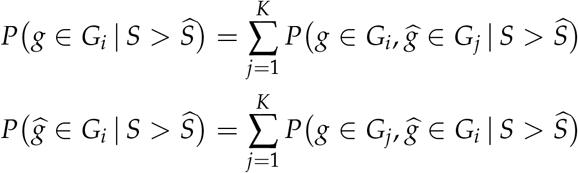

to obtain

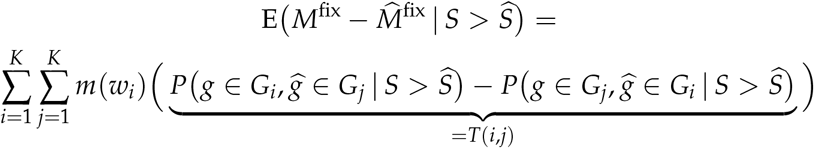

Consider the inner terms of the sum, which we denote by *T*(*i, j*). Notice that *T*(*i, i*) = 0 and *T*(*i, j*) = −*T*(*j, i*). Because of this

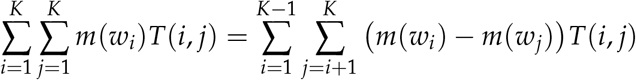

which written out becomes

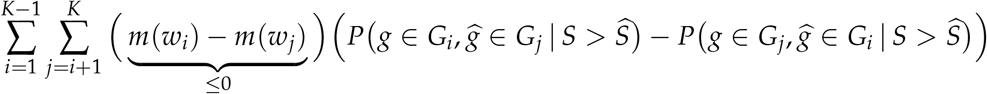

When *i* = 1, there is at least one *j*_0_ in {2, ⋯ *K*} such that *m*(*w*_1_) – *m*(*w*_*j*_0__) < 0 by assumption. We now look at the conditional probabilities in the product. We have

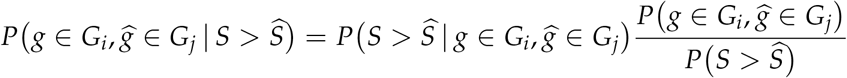

Because *G* and 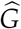 are independent and identically distributed, we have

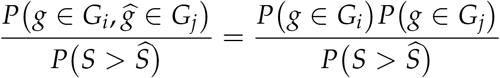

We therefore get

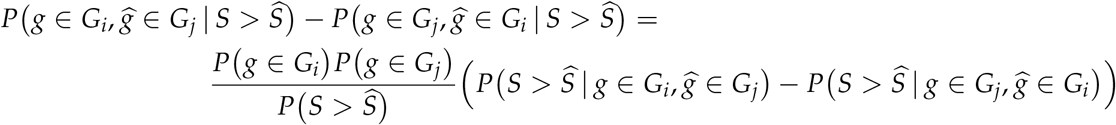

Looking at the second term of the difference, because of symmetry and independence, we can write

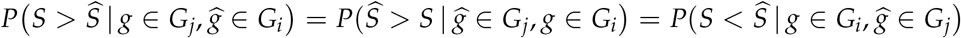

Now it is time to use our assumption

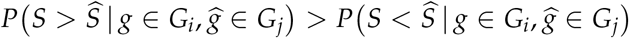

to conclude that the difference is positive

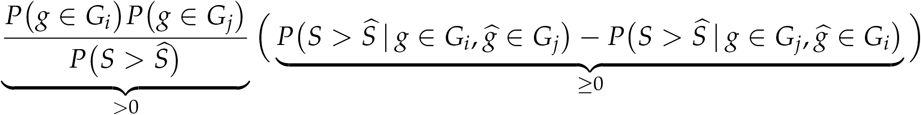

with the inequality being strict at least when *i* = 1 and *j* = *j*_0_, since *w*_1_ <*w*_*j*_0__. It follows that *dM* < 0

#### The case of positive selection

The case of positive selection is entirely analogous to the case of negative selection presented above. The only modification is that for the fitnesses we now have *w*_1_ > *w*_weak_, and the stochastic dominance assumption is modified as:

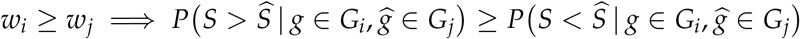

The proof proceeds in an identical manner, with changes in the appropriate signs, to yield *dM* > 0.

### A test statistic for selection

The quantity we use as basis for our test is

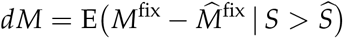

We have shown that *dM* = 0 if there is no selection, whereas *dM* < 0 under negative selection and *dM* > 0 under positive selection. We again note that it is possible that *dM* = 0 even if we have selection (details above), and in this sense the test we propose will only have power under some alternatives: if we observe *dM* ≠ 0 we can confidently conclude selection whereas *dM* = 0 is compatible with both no selection and - if certain assumptions are not met - selection.

Now, suppose we sample an individual from the population. *dM* is defined based on an expectation over possible evolutionary trajectories and our observed data only represent the realization of one of these trajectories. To circumvent this, we leverage the fact that we observe the epigenetic state throughout the entire genome. Our test statistic is an empirical approximation of *dM* defined as

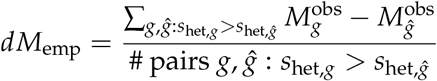

where 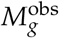 is the observed epigenetic state at gene *g* (equal to 1 is *M* is observed and 0 otherwise). Each term in the sum has values in {−1, 0, 1}.

This test is essentially testing for stochastic monotonicity between *M* and *s*_het_. Hence, to obtain the distribution of the test statistic under the null hypothesis of no selection, we use permutations. Specifically, we permute the selection coefficients associated with each gene: this reflects the null hypothesis because without selection there is no association between the fixation probability and genic *s*_het_.

For visualization, we use the following quantity (or a variant thereof): we bin genes into deciles based on their *s*_het_. For each decile, we plot the proportion of genes with the *M* epigenetic state, that is

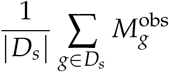

where *D_s_* is a given decile. Under the scenario of no selection, the resulting curve will be flat (with random fluctuations), whereas if there is selection the curve will be increasing or decreasing.

Finally, we explicitly state two approximations that are implicit in our test. For the first approximation, we treat the population-scaled “epimutation” rate as being low enough, so that an epigenetic state introduced into the population at a given gene always undergoes fixation or extinction before a new epigenetic state arises at the same gene. Based on this, we consider a randomly sampled individual from the population as having the resident epigenetic state at the vast majority of genes (we note that a similar assumption underlies the popular 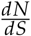 test, see e.g. Nielsen, Yang (2003) and Kryazhimskiy, Plotkin (2008)). In the case of proximal promoter DNA methylation, we performed an analysis of changes along primates and found that the genome-wide epimutation rate across promoters is in fact lower than the coding mutation rate (Supplemental Figure S10; see also section “Promoter DNA methylation “epimutation” rate”), providing support for this approximation. In addition, we further examined the validity of this approximation by comparing the nucleotide diversity of methylated CpGs (+/− 2kb from the TSS) to the nucleotide diversity of hypomethylated CpGs in the same regions, as estimated from the sequences of 62,874 individuals (Taliun et al. (2021); see section “Genetic Variation data from TOPMed”). We found that the methylated CpGs have substantially higher diversity (Supplemental Figure S11). This confirms that, while the methylation state was determined based on samples from only 2 individuals (Molaro et al., 2011), these 2 individuals suffice to obtain an accurate picture of the methylation state of the majority of the population. The second approximation we make is that in our definition of *dM*_emp_ we view each gene pair as an independent realization of the evolutionary process; this is only true for genes belonging to different regulatory groups.

### Epigenetic marks with continuous states

The framework above can readily accommodate epigenetic marks with continuous states. In that case, we no longer have just two different states *U* and *M*. Instead, we have a mark whose state, denoted by *M*, attains values in a continuous interval. Then, the only difference is that 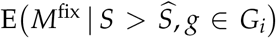 corresponds to the expected value of the state at equilibrium, instead of the fixation probability. This again depends only on the fitness effect of *M* at group *i*, and can be written as *m*(*w_i_*). The proof is otherwise identical.

We note that in some cases, it may be possible to obtain the continuous behavior of an epigenetic state as the sum of latent binary states. As an example, let us consider the case of the size of the hypomethylated region around hypomethylated proximal promoters. Let the total region around a promoter consist of *N* sub-regions, each of which can exist in a methylated or hypomethylated state. Let *M*_total_ be the (continuous) stochastic variable representing the size of the methylated region and let *M_i_, i* = 1,…, *N* be a set of binary stochastic variables as above, with 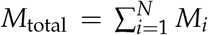. Then we have

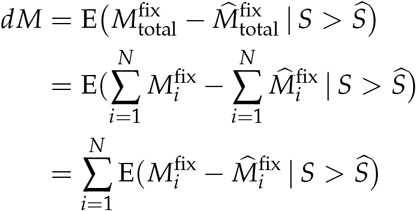

While we do not investigate this further, we mention that the observed states of other marks may be thought to arise in a similar fashion. For example, the ChIP-seq signal of a histone modification at a given region may be viewed approximately as the additive contribution from the different nucleosomes present in that region.

For visualization, similar to binary marks, we first bin genes into deciles. For each decile, we then plot the distribution of the states.

### Adjusting for the effect of confounding molecular features

It is important to test if the observed signal of selection on a given mark is in fact a spurious signal arising as a passive consequence of selection on a different molecular feature. To address this, we consider a counterfactual *dM*:

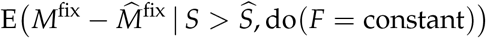

where *F* stands for any other molecular feature we wish to test, and do (*F* = constant) means that the value of *F* is fixed to the same level for every gene in the genome.

If *F* is entirely responsible for the selection signal on *M*, then

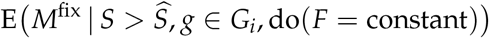

will be the same for all groups. This is because the underlying variation that drives differences in the fixation probability (or, for epigenetic marks with continuous states, the expected value at equilibrium) between different groups is eliminated. Therefore, as in the scenario of neutrality, the counterfactual *dM* will be equal to 0.

The rationale behind this counterfactual *dM* becomes clear by recognizing the analogy between our setup for selection inference, and standard causal inference (Imbens, Rubin, 2015). In our case, the “unit” is the population of individuals, a certain proportion of which have the epigenetic state associated with a gene of interest. The “exposure” is the genic *s*_het_, and the “potential outcome” is the fixation of the epigenetic state. Note that in our case, the potential outcome is inherently stochastic due to the action of random genetic drift; this is a difference with the traditional causal inference framework, where the potential outcome is treated as a deterministic function of the exposure (Imbens, Rubin, 2015).

In the case of a spurious signal, selection acts on the intermediate feature. But because the intermediate feature causally determines the epigenetic state, our test yields a result consistent with selection on that state. In other words, the effect of the exposure on the fixation of the epigenetic state is mediated through the exposure’s effect on the fixation of the intermediate feature. Using terminology from mediation analysis (Pearl, 2014; VanderWeele, 2016), we want to know if there is a statistically significant controlled direct effect of *s*_het_ on the fixation probability of the epigenetic state (in other words, whether or not there is complete mediation).

Traditionally in mediation analysis, this is achieved fitting a regression model with the outcome as the response, and testing if the partial regression coefficient of the exposure while adjusting for the mediator is different than zero (James et al., 2006; MacKinnon, 2012; VanderWeele, 2016). In the case of linear regression, this is mathematically equivalent to a residual-on-residual regression, with the two residuals obtained by regressing the outcome on the mediator and the exposure on the mediator, respectively. Following this, we perform our test as follows. If the epigenetic state is binary, we first fit a logistic regression model across genes using the observed state as a binary response and the feature we wish to adjust for as a covariate. We then obtain the deviance residuals from this model, which are bimodal, and treat the binarized version of these residuals as the adjusted version of the binary state. If the epigenetic state is continuous, we instead use the residuals from a linear regression of the state on the confounding feature. In both the binary and the continuous case, we then apply our test on the adjusted epigenetic state, but instead of *s*_het_ we use the residuals from a linear regression of *s*_het_ on the feature.

As with all applications of mediation analysis, it is important to be aware that if the observations of the mediator are too noisy, the presence of a controlled direct effect may be inferred, even when the true causal model is one of complete mediation (Ledgerwood, Shrout, 2011; Otter et al., 2018; Gastonguay et al., 2022). In our framework, this means that if the intermediate, potentially confounding feature (e.g. gene expression) is measured with too much noise, one can erroneously infer that the selection on the epigenetic state is not a passive byproduct of selection on the intermediate feature, even when the opposite is true. We further investigated this by simulating simple idealized scenarios with linear mediation and measurement corrupted by normally distributed noise, and performed 10000 simulations under complete mediation for each of several levels of noise. We found that, as long as the variance of the noise is less than 10% of the variance of the signal, type I error control in the inference of a controlled direct effect is maintained. Notwithstanding this, the results from the analysis of potential mediation driving selection should generally be corroborated by orthogonal lines of evidence. In our case, they are consistent with targeted epigenome editing experiments (see main results section and references therein).

### A model for the interaction between an epigenetic state and a coding variant

Finally, given that we are focusing on epigenetic marks associated with genes, we briefly present a simple model for interactions between epigenetic states and coding variants in the associated gene. We note that this is not used in our selection test.

First, consider a single gene with a biallelic site with coding alleles *A* and *D* in the population. Let *D* correspond to a loss-of-function allele, with the selection coefficient against heterozygotes equal to *s*_het_. The fitness *w* is then given by

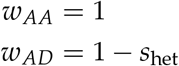

with 0 ≤ *s*_het_ ≤ 1. We ignore the *DD* genotype, since selection happens predominantly through heterozygotes (Falconer, MacKay, 1995; Fuller et al., 2019).

We extend this model with an additional site in the same gene, where an epigenetic mark is present, with possible epigenetic states *U* and *M*. Since *D* is a loss-of-function allele, we will assume that the fitness effect of the *D*, *U* and *D*, *M* combinations is the same when *D* and *U*/*M* reside on the same chromosome, and just use *D* to designate them (in other words, if a coding loss-of-function allele is present, the epigenetic state has no extra fitness effect). We then write the fitness now as:

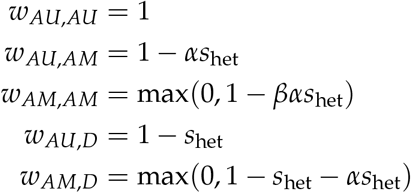

with *β* ≥ 1.

### Data processing and analysis

#### Promoter coordinates and transcript selection

We defined proximal promoters as 1kb regions centered around the transcriptional start site (TSS). We obtained a set of 11,059 promoters with high-confidence GENCODE TSS annotation provided in Boukas et al. (2020a). As described therein, this set does not contain subtelomeric promoters (within 2 Mb of chromosome ends), as the CpG islands of such promoters have distinct characteristics (they are organized in clusters, and are thought to be maintained principally by GC-biased gene conversion (Cohen et al., 2011)). Also excluded are promoters of genes on the sex chromosomes, for which loss-of-function intolerance estimates have a different interpretation than for autosomal genes, owing to hemizygosity in males/X inactivation in females.

We further restricted to promoters where the downstream transcript had ≥ 10 expected loss-of-function variants, in order to ensure that our rank ordering of genes according to the selective pressure against heterozygous loss-of-function alleles is not severely corrupted by genes not adequately powered for LOEUF or *s*_het_ estimation. Subsequently, bidirectional promoters were handled exactly as described in Boukas et al. (2020a), yielding a set of 7,518 promoters that we used as input for our downstream analyses.

To exclude the possibility that our stringent filters for TSS selection led to a biased result, we tested for selection on proximal promoter DNA methylation, this time using CAGE data from the FANTOM5 project to define promoter coordinates. Since our focus is on DNA methylation in the male germline, we used data from human testis (http://fantom.gsc.riken.jp/5/datafiles/latest/basic/human.tissue.hCAGE/testis%252c%2520adult%252c%2520pool1.CNhs10632.10026-101D8.hg19.ctss.bed.gz). This bed file contains TSS coordinates as defined by CAGE-seq. We grouped the TSS’s by gene, and for each gene chose the TSS corresponding to the transcript with lower LOEUF. If a gene still had more than one TSS, we chose the TSS with the highest value in the “score” column. For genes that still did not have a unique TSS, we randomly selected one of the TSS’s. These approach allowed us to study 12,967 promoters, a substantial increase compared to the 7,235 promoters we studied before (note that there are still several promoters that are excluded because we require the downstream transcript to have ≥ 10 expected loss-of-function variants). Reassuringly, the results with these FANTOM5-based promoter coordinates were the same. We found strong evidence for selection against the presence of DNA methylation at the proximal promoter; *dM* is equal to −0.08 (*p* = 9.9 · 10^-4^), very close to our original *dM* of −0.07.

Finally, we also performed our test of selection on proximal promoter DNA methylation using only the portion of the promoter upstream of the TSS. That is, we do not include any 5′ UTR or coding sequence. For this analysis, we defined the upstream promoter portion as 750bp upstream of the TSS, to ensure it includes enough CpGs. The results were in full agreement with our original results, with negative selection against proximal promoter DNA methylation (*dM* = −0.04; *p* = 9.9 · 10^-4^).

#### Whole-genome bisulfite sequencing data from human sperm

We used processed whole-genome bisulfite sequencing data from human sperm (Qu et al., 2018). These data were accessed though the DNA methylation trackhub at the UCSC genome browser (hg19 assembly; Song et al. (2013)), and consisted of methylation level (defined as the proportion of reads supporting the methylated state) and coverage. The raw experimental data consisted of two biological replicates (Molaro et al., 2011). We note here that we chose sperm because it has been established that the majority of CpG>TpG transitions occur in the male germline. For example, a recent analysis of trio genomic sequencing data showed that the distribution of *de novo* CpG>TpG transitions along the genome is strongly correlated with genome-wide methylation levels in testis, and much less so with methylation levels in ovaries (Gao et al., 2019). This implies that a major fraction of these mutations arise after germ cell specification, and not during the period from zygote until germ cell formation, where the methylation landscape of both male and female cells is similar. Further underscoring the major contribution of male germline methylation to the load of CpG>TpG mutations, these mutations show a 6.5-fold yearly increase in males compared to females following puberty (Gao et al., 2019).

We only considered CpGs with at least 10× coverage, and restricted to proximal promoters with ≥ 10 CpGs (7,235 promoters in total). To compute the percentage of methylated CpGs in each proximal promoter, we labeled a given CpG as methylated if its methylation level was ≥ 80%, and hypomethylated if its methylation level was ≤20%; CpGs with intermediate methylation level (that is, between 20% and 80%) were discarded. As orthogonal support for the methylation state of the human promoter CpG sites, we examined their nucleotide diversity and minor allele frequency spectrum in TOPMed (see section “Genetic variation data from TOPMed” below); reassuringly, methylated CpGs are substantially more variable than hypomethylated ones (Supplemental Figure S11a, b).

To ensure that the result of our test on selection on proximal promoter methylation is not driven by the thresholds used to define the methylation status of the CpGs, we also calculated the average methylation (across all CpGs) for each proximal promoter. We then applied our test again, this time treating this per-promoter average methylation as a continuous mark, and obtained very similar results (*dM* = −0.06; *p* = 9.9 · 10^−3^ after 1000 permutations).

#### Hypomethylated regions

To obtain the coordinates of hypomethylated regions in human sperm – as detected in Qu et al. (2018) – we first downloaded the corresponding BigBed (“.bb”) file from the DNA methylation trackhub at the UCSC genome browser (hg19 assembly). We then converted this BigBed file into a BED file, and imported it into R using the “rtracklayer” package.

When assessing the relationship between hypomethylated region width and LOEUF, we sought to minimize confounding by hypomethylated regions that are large because they correspond to bidirectional promoters. To this end, we adopted a more stringent procedure than the one used to exclude bidirectional promoters in Boukas et al. (2020a). Specifically, out of the high-confidence promoters we used for the analysis of proximal promoter methylation, we further excluded promoters which overlapped regions +/− 4kb from any TSS corresponding to a transcript. This was done after obtaining coordinates of such TSSs with the promoters() function from the “EnsDb.Hsapiens.v75” R package. Our procedure resulted in 3,580 transcripts.

#### Transcriptional end regions

We obtained coordinates of transcriptional end sites using the transcripts() function from the “EnsDb.Hsapiens.v75” R package. We defined transcriptional end regions as regions +/− 500bp on either side of the transcriptional end site. We estimated the methylation state of these transcriptional end regions by following the exact same steps as for proximal promoters. To avoid the detection of a spurious selection signal driven by selection on promoter methylation, we excluded transcriptional end regions that overlapped regions +/− 2kb from any TSS corresponding to a transcript (coding or non-coding). We then also restricted to transcripts with at least 10 expected loss-of-function variants in gnomAD. These filters resulted in 4,545 transcriptional end regions. Finally, as with proximal promoters, we also tested for selection by computing the average methylation (across all CpGs) of each end region, and treating it as a continuous mark. This yielded very similar results (*dM* = 9.54 · 10^−5^; *p* = 0.487)

#### H3K36me3 ChIP-seq data

We downloaded the raw ChIP-seq reads (fastq files) from GSE40195. Reads were mapped to hg19 using Bowtie2 (Langmead, Salzberg, 2012). Subsequently, we used Picard (http://broadinstitute.github.io/picard/) to remove duplicate reads, with the function MarkDuplicates. We then called broad peaks using MACS2 (Y Zhang et al., 2008), with the “keep-dup” parameter equal to “all”. We chose to perform our main analyses using these broad peaks, as H3K36me3 is a mark that tends to show broad distribution. However, to ensure that the result of positive selection on coding H3K36me3 is not dependent on this choice, we also performed our test after calling narrow peaks and obtained a very similar result (*dM* = 0.07, *p* = 9.9 · 10^−4^).

To investigate the relationship between genic loss-of-function intolerance and exonic H3K36me3, we first obtained coordinates of coding exons using the cdsBy() function from the “EnsDb.Hsapiens.v75” R package. We then restricted to canonical transcripts (as provided by gnomAD; 18,912 transcripts total), and identified all such transcripts that have at least one H3K36me3 peak in their coding sequence using the findOverlaps() function from the “GenomicRanges” R package. We then restricted to transcripts that have at least 10 expected loss-of-function variants in gnomAD. Ultimately, we examined 13,361 transcripts.

#### H3K4me3 ChIP-seq data

We used the “AnnotationHub” R package to obtain processed H3K4me3 data from human H1 embryonic stem cells (sample ID “UCSD.H1.H3K4me3.LL312”). These data include the locations (genomic coordinates) of narrow peaks, as well as the intensity of the signal at each peak (column named “signalValue”). If a given promoter harbored more than one peak, we computed the promoter H3K4me3 signal as the average of the signals for each peak.

#### Permutation p-values

We calculated permutation p-values following Phipson, Smyth (2010), with the number of permutations being either 10,000 or 1,000.

#### Triplosensitivity estimates

We obtained probability of triplosensitivity (pTS) values for human genes from the supplemental material of Collins et al. (2021). Since this supplemental material provides only gene names as the identifier, we matched these gene names to transcript ids using the “EnsDb.Hsapiens.v75” R package.

#### eQTL datasets

We obtained eQTLs corresponding to testis from the GTEx portal (GTEx Consortium,2020) and eQTLs corresponding to human induced pluripotent stem cells from the supplemental material of DeBoever et al. (2017). These datasets provide information about eQTLs mapped using genotype data from 322 and 215 individuals, respectively. This information includes the effect size and p-value corresponding to each variant. For testis, following standard GTEx practices, we used the allelic fold-change (in *log*_2_ scale) as the measure of effect size. For induced pluripotent stem cells, since the allelic fold change was not provided, we used the estimated betas.

#### Gene expression data

For expression in the male germline, we downloaded the gene-level TPM expression values in testis from the GTEx v7 release (GTEx Consortium et al., 2017), from the GTEx portal. We restricted to the genes downstream of our 7,518 promoters. Subsequently, for each gene we computed the median (across individuals) expression in testis (in log_2_ (*TPM* + 1) scale).

We expression in embryonic stem cells, we downloaded the raw fastq files from GSE90225, corresponding to RNA-seq data in H1 embryonic stem cells generated by ENCODE (ENCODE Project Consortium, 2012). We pseudoaligned these reads to a fasta file containing all human cDNA sequences (Homo_sapiens.GRCh37.75.cdna.all.fa; downloaded from ENSEMBL), and subsequently quantified transcript abundances with Salmon (version 0.10.0) (Patro et al., 2017). We then obtained normalized gene-level counts from the transcript abundances using the tximport R package (Soneson et al., 2015), with the “countsFromAbundance” parameter equal to “lengthScaledTPM”. We subsequently performed all analyses in log_2_ (*TPM* + 1) scale.

For fetal gene expression data, we downloaded the “Proportion Matrix by Cell Type” from https://descartes.brotmanbaty.org/bbi/human-gene-expression-during-development/. This matrix contains the proportion of cells expressing a given gene (UMI > 0) in each of 172 fetal cell types identified with single-cell RNA-seq (Cao et al., 2020).

#### Identification of genes whose promoters change methylation status between the germline and embryonic stem cells

We downloaded whole-genome bisulfite sequencing data corresponding to human h9 embryonic stem cells in an identical manner with the sperm data. We then calculated the sperm and H9-ESC proximal promoter methylation of all canonical transcripts, and selected the proximal promoters that are hypomethylated in H9-ESCs (≤20% CpGs methylated) but methylated in sperm (≥ 80% CpGs methylated). We subsequently sought to exclude genes whose expression in the germline may be driven by some alternative hypomethylated promoter, which then also potentially drives expression in embryonic stem cells. To this end, we restricted to genes whose expression in the germline (testis; GTEx Consortium et al. (2017)) is less than 1 (in log_2_ (*TPM* + 1) scale). Finally, to match the embryonic stem cell expression between genes whose proximal promoters are methylated in the germline but become hypomethylated in ESCs, and gene whose proximal promoters are hypomethylated both in the germline and ESCs, from the latter group we excluded genes with expression greater than 8.

#### Association between coding mutation rate and loss-of-function intolerance

We downloaded all synonymous *de novo* mutations from the gene4denovo database (total of 15,642 mutations; Zhao et al. (2020)). For each transcript, we normalized the number of synonymous mutations by the total length of the coding sequence, and subsequently restricted to transcripts with at least 10 expected loss-of-function variants in gnomAD (13,361 transcripts in total). However, the assessment of the relationship between coding *de novo* mutation rate and loss-of-function intolerance is complicated by the fact that there are genes which have no *de novo* mutations in the database. The interpretation for these genes differs according to the length of their coding sequence (observing 0 mutations implies a different mutation rate in a gene whose coding sequence length is 3kb compared to a gene whose coding sequence length is 1kb). To address this issue, we performed the following analyses. First (corresponding to Figure 4), we excluded genes with 0 mutations. Second, we included all genes but added a pseudocount of 1 to each gene; the results were highly similar (Supplemental Figure S9a, b).

As an orthogonal measure of mutation rate, we used the synonymous substitution rate (dS), calculated from a comparative analysis of: a) human vs chimp, and b) human vs mouse. We obtained these dS values from ENSEMBL version 96 using the biomaRt R package, and then assessed the relationship between LOEUF and dS after excluding genes with dS equal to 0. Again, we found that greater loss-of-function intolerance is associated with lower mutation rate (Supplemental Figure S9c, d).

We note here that, after our preprint was posted, another study reported no association between *de novo* mutation rate and genic selective constraint in humans (H Liu, J Zhang, 2022). However, that report examined a much smaller dataset of *de novo* mutations. Thus, due to the paucity of data, they were forced to calculate mutation rates over the entire gene sequence, after including not only synonymous mutations, but also intronic mutations, as well as missense and loss-of-function ones. Prompted by the results of H Liu, J Zhang (2022), we examined how the choice of the specific types of *de novo* mutations used to calculate the mutation rate impacts the results. Using our (much larger) dataset, we found that the inclusion of intronic mutations leads to a big reduction in the strength of the association between LOEUF and *de novo* mutation rate (Supplemental Figure S9e,g,h; adjusted *r*^2^ = 0.7 when including intronic mutations vs 3.8 with synonymous mutations only). This emphasizes that the effect is highly localized to the regions under selection: coding exons. This locality also supports the notion that the effect is driven by features like epigenetic marks. Finally, if one also includes non-synonymous coding mutations in addition to intronic ones, the association disappears (Supplemental Figure S9g,h). This is most likely attributed to the fact that a lot of the studies on *de novo* mutations include sequencing data from individuals with Autism Spectrum Disorder and developmental disorders, and are thus enriched for damaging mutations in loss-of-function intolerant genes.

#### Between species nucleotide conservation

We quantified nucleotide conservation across 100 vertebrates species with the PhyloP score (Pollard et al., 2010). We obtained these scores for nucleotides in promoters with the phastCons100way.UCSC.hg19 R package.

#### Estimation of promoter CpG mutation rates

We first obtained processed whole-genome bisulfite sequencing data from chimp and rhesus sperm (Qu et al., 2018) in an identical manner to the human data. Briefly, these data were generated as follows. First, reads were mapped to the corresponding chimp and rhesus assemblies to generate methylation and coverage level, and the CpGs were subsequently aligned to their homologous position in hg19 (which need not be a CpG; see Qu et al. (2018) for details). We restricted to CpGs which had more than 10× coverage and overlapped human proximal promoters. In each species, we categorized CpGs whose methylation level was ≥ 80% as methylated, and CpGs whose methylation level was ≤20% as hypomethylated.

We restricted to CpGs in promoters within the top LOEUF quartile, in order to minimize the effect of selection on our mutation rate estimates. Using rhesus as the outgroup, we subsequently defined ancestrally methylated CpGs as those methylated in both chimp and rhesus (1,490 CpGs), and ancestrally hypomethylated CpGs as those hypomethylated in both chimp and rhesus (42,732 CpGs). We found that 118 of the methylated and 1,090 of the hypomethylated CpGs, respectively, are mutated in human. Using a generation time of 25 years for chimp, and a human-chimp divergence time of 8 million years, this yielded mutation rate estimates of 2.47 · 10^−7^ for methylated CpGs and 7.97 · 10^−8^ for hypomethylated CpGs.

Across the entire genome, approximately 65% of CpGs are hypomethylated and 14% are methylated in the male germline. Based on these percentages, and assuming that the genome-wide CpG mutation rates are similar to our promoter CpG mutation rates, our estimates are consistent with the genome-wide estimate obtained by parent-offspring trio sequencing in (Kong et al., 2012). Moreover, the 3.10-fold increase in mutation rate conferred by methylation is concordant with estimates recently provided in Zhou et al. (2020).

#### Estimation of exonic mutation rates within and outside H3K36me3-marked regions

To estimate these mutation rates, we first calculated the ratio of the number of synonymous mutations per base pair in non-H3K36me3-marked exonic sequences to the corresponding number for H3K36me3-marked exonic sequences (+/− 250 bp on either side of an H3k36me3 peak excluding intronic sequence) as 1.23. Then, combining the marginal exome-wide mutation rate estimate of 1.38 · 10^−8^ per site per generation from Rodriguez-Galindo et al. (2020), with the fact that H3K36me3-marked exonic sequences form 8% of all exonic sequences, we arrived at the system of equations

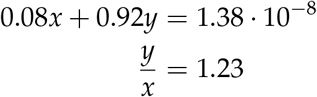

where *x* and *y* are the mutation rates of H3K36me3-marked and non-H3K36me3-marked exonic regions, respectively. Solving the system yields values of 1.14 · 10^-8^ for *x* and 1.4 · 10^-8^ for *y*.

#### Estimation of promoter DNA methylation “epimutation” rates

Using whole-genome bisulfite sequencing data from chimp and rhesus sperm (see “Estimation of promoter CpG mutation rate” section above) we categorized human proximal promoters as methylated (≥ 80% methylated CpGs) or hypomethylated (≤20% methylated CpGs) in chimp and rhesus.

To minimize the influence of selection on our estimates, we again focused on promoters within the top LOEUF quartile. Using rhesus as the outgroup, we then defined ancestrally methylated promoters as those methylated in both chimp and rhesus (387 promoters), and ancestrally hypomethylated promoters as those hypomethylated in both chimp and rhesus (1301 promoters). We found that 4 of the methylated and 1 of the hypomethylated promoters, respectively, have the opposite methylation status in human. Using the same estimates for chimp generation time and human-chimp divergence as above, this corresponds to “epimutation” rate estimates of 3.23 · 10^-8^ for methylated promoters and 2.4 · 10^-9^ for hypomethylated promoters.

#### Genetic variation data from TOPMed

We downloaded a VCF file containing human variation data from dbSNP (version 151, hg19 assembly). We then used bedtools (Quinlan, Hall, 2010) to restrict to variants within our set of 7,518 promoter regions, extended to 2kb on either side of the TSS. We used the allele frequencies of these variants in TOPMed (freeze 5; 62,874 individuals), and only considered biallelic sites with single nucleotide variants (that is, we excluded multiallelic sites and sites with indels). We further restricted to sites where the reference allele was the major allele (allele frequency ≥ 0.5). These filters resulted in a total of 4,568,818 sites, of which 707,637 were CpGs. Following Asthana et al. (2007), the nucleotide diversity (*π*) for a given set of sites was estimated as

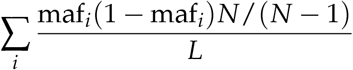

where *L* is the total number of sites, maf_*i*_; is the minor allele frequency in site *i*, and N is the total number of individual chromosomes (which in our case is equal to 125,748 as there are 62,874 individuals). For sites with no variant allele, the minor allele frequency was taken to be 0.

### Selection on regional epigenetic mutation rate modifiers

#### Infinite population size

In this and the following section, we denote the epigenetic states as A and B instead of U and M, in order to maintain terminology consistent with the standard population genetics literature.

Let us consider an effectively infinite haploid population consisting of individuals of type A and B, corresponding to the two states of the epigenetic mark of interest. The time evolution of the population is described by the equations

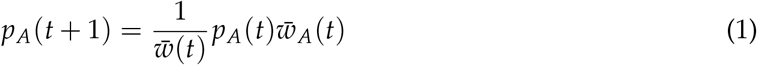

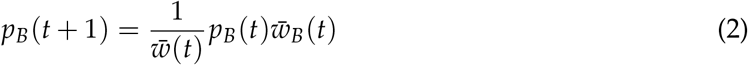

where the average fitnesses of type *A* and type *B* individuals are given by 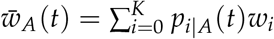 and 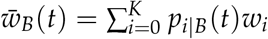, respectively, and 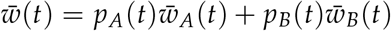.

To model the conditional distributions *p*_*i*|*A*_ and *p*_*i*|*B*_, we make the following assumptions:

- The number of new mutations harbored by an individual after each generation follows a Poisson distribution with rate parameter equal to *U_A_* and *U_B_* for the two types, respectively
- Each mutation has the same selection coefficient *s*
- The relative fitness of an individual with *i* mutations is given by *w_i_* = (1 – *s*)^*i*^
- Each new mutation arises at a unique site
- There are no backmutations
- There is no recombination. We note that recombination can play a very important role in the case of genetic mutation rate modifiers (such as DNA repair gene variants), because it can break the association between the mutator allele and the mutation-loaded genome, thus preventing further accumulation of mutations at an elevated rate. However, epigenetic mutation rate modifiers are not subject to recombination themselves. Recombination may temporarily reduce the genomic mutation load, but mutations will continue to accumulate at an elevated rate.

Then, the process of mutation accumulation gives rise to the following time dynamics for *p_i|A_* and *p_i|B_* (Kimura, Maruyama (1966), Haigh (1978), see also Maia et al. (2003) for an explicit exposition):

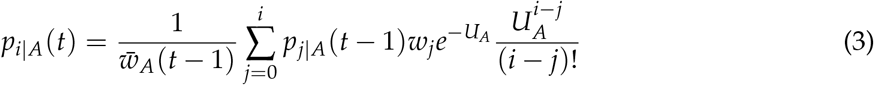

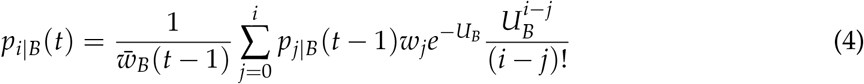

At within-type equilibrium, i.e. when *p_i|A_*(*t*) = *p_i|A_* (*t* – 1) and *p_i|B_* (*t*) = *p_i|B_* (*t* –1), it is a standard result (Kimura, Maruyama, 1966) that 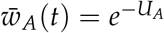 and 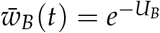. From equations (1) and (2) it then follows that, for all t after within-type equilibrium has been reached:

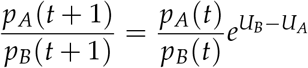

But *U_B_* > *U_A_*, so *e*^*U_B_* – *U_A_*^ >1. Therefore,

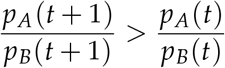

#### Finite population size

We simulated a population with effective size *N* = 1000, consisting of two types of individuals. As before, type A corresponded to individuals with the *U* epigenetic state, and type B to individuals with the *M* epigenetic state. Each region consisted of 80 sites for the simulations corresponding to proximal promoter DNA methylation, and 135 sites for the simulations corresponding to exonic H3K36me3; these numbers were chosen based on the relevant genome-wide averages. We note that the choice of 1000 for the effective population size was made to strike a compromise between computational efficiency and the actual value for vertebrates (10^4^). To match this, we also scaled the mutation rate estimates by multiplying by 10; we verified that these scalings do not affect the results.

At *t* = 0, we assumed 500 individuals of type A and 500 individuals of type B, and that all individuals of both types harbored 0 mutations. Then, each generation was generated by the previous one assuming the following life-cycle.

- First, within each type, mutation and selection acted in an infinite population of gametes, as described in equations (3) and (4) of the previous section.
- For computational efficiency, we viewed 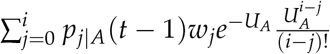 as the convolution of *x*[*n*] = *p*_*n|A*_ (*t* – 1)*w_n_* and 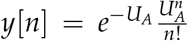. We computed the convolution using the Fast Fourier Transform. Specifically, we used the “fftwtools” R package (Rahim,2021), which provides the “Fastest Fourier Transform in the West” implementation (Frigo, Johnson, 2005).
- Then, from the resulting, still infinite, population of gametes, a sample of 1000 gametes were allowed to survive to “adulthood” and form the next generation of individuals. These 1000 gametes were chosen via multinomial sampling, with the probability of choosing a gamete with *i* mutations given by

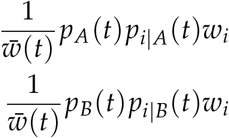

for gametes of type A and type B, respectively. This gave the next generation of individuals, from which the updated type A and type B frequencies, as well as the updated type-conditional frequencies of individuals with *i* mutations, were calculated.

We investigated different scenarios, corresponding to simultaneous upstream control of the epigenetic state of different numbers of locations (Figure 6). In each scenario, we allowed the product *Ns* to obtain different values. Specifically, we used values at the border of effective neutrality for CpGs (Figure 6), based on the results of Kryukov et al. (2005) for conserved non-coding sites. By contrast, we used higher selection coefficients for coding sites (Figure 6, Kryukov et al. (2005)). For each of these sub-scenarios, we performed 150 forward simulations, and each simulation lasted until either type A or type B reached fixation.

#### Diffusion approximation to the fixation probability of a mutation rate modifier affecting a small number of sites

For notational convenience, in this section we move *t* to the subscript. Let *U* (*p_B,t_*) be the probability that the *M* state reaches fixation, conditional on its current frequency at time *t* being equal to *p_B,t_*, and conditional on the fitnesses *w_A,t_* and *w_B,t_*. It is well-known that *U* satisfies the Kolmogorov backward equation:

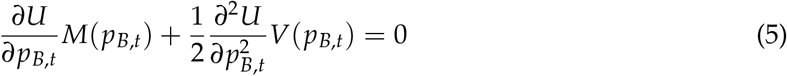

where *M* is the conditional expectation of the change in allele frequency from the present generation to the next, and *V* is the sum of *M*^2^ and the conditional variance of the change in allele frequency from the present generation to the next. Both *M* and *V* are conditional on the present allele frequency *p_B,t_*, and on *w_A,t_* and *w_B,t_*.

We have:

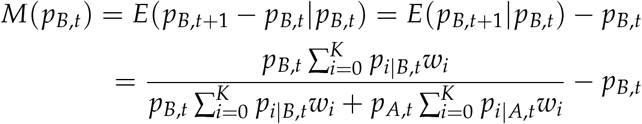

Now let us consider the situation where *M* causes only a negligible increase in the mutation rate, which is the case when it affects a small number of sites. In that case, we expect that, on average, the conditional distributions *p_i|A_* and *p_i|B_* would be the same. Therefore, we have:

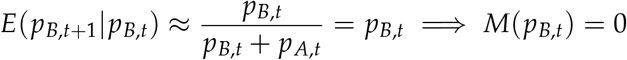

And,

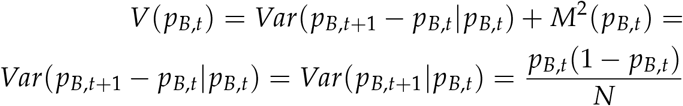

where *N* is the effective population size.

From Equation (5), since *V* (*p_B,t_*) ≠ 0, we obtain

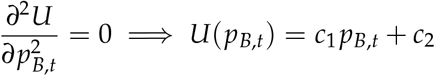

where *c*_1_ and *c*_2_ are constants.

Using the fact that *U*(0) = 0 and *U*(1) = 1, we arrive at *U*(*p_B,t_*) = *p_B,t_*.

This is what we would intuitively expect to be true: when the modifier only affects a small number of sites, and thus causes a negligibly small increase in the total mutation rate, the probability of fixation of the modifier is equal to its present frequency, which is the result under neutrality.

## Code availability

The code used for the analyses and figures is available at https://github.com/hansenlab/epigenetics_selection.

## SUPPLEMENTARY MATERIALS

### Supplemental Figures

**Supplementary Figure S1.**
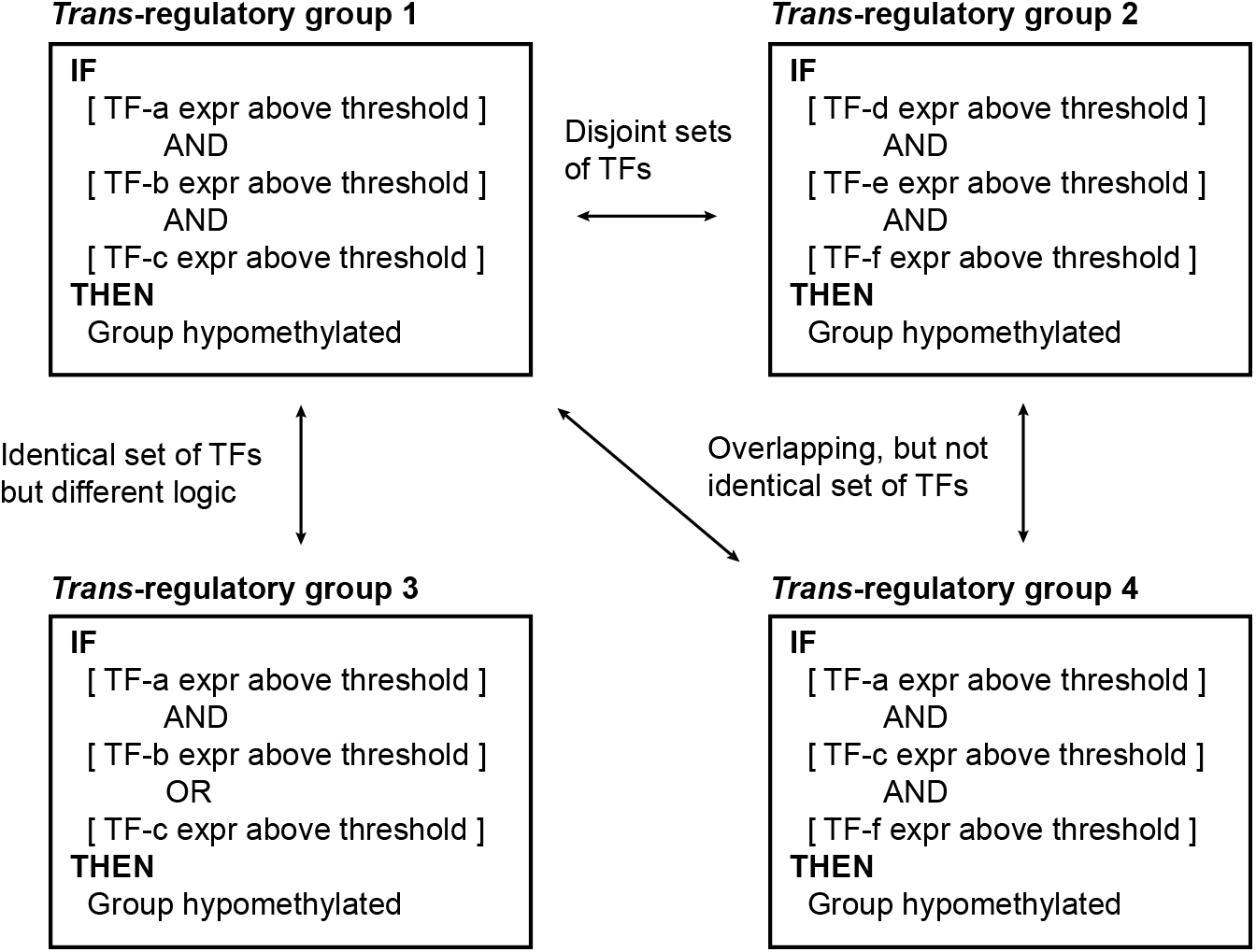
Examples of different *trans*-regulatory groups. Four different hypothetical *trans*-regulatory groups defined by the logic based on which the corresponding regulatory factors interact to determine the epigenetic state (here, DNA methylation states).

**Supplementary Figure S2.**
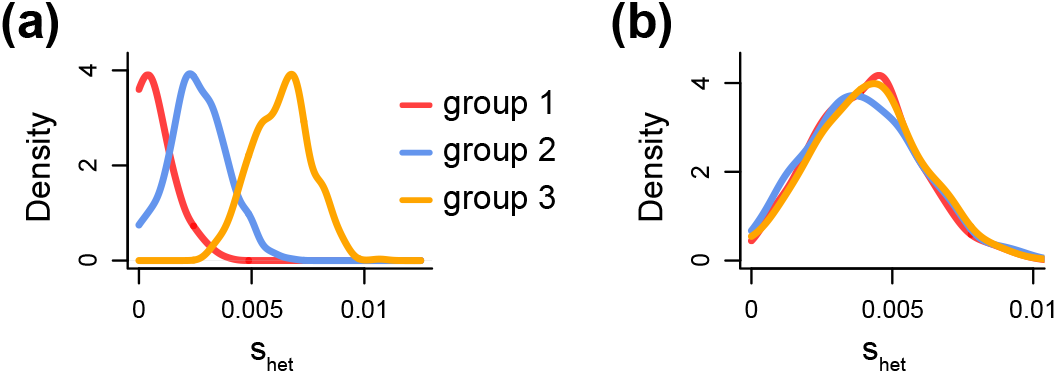
Hypothetical *s*_het_ distributions of 3 different *trans*-regulatory groups. We assume that for the fitness effect of the epigenetic mark at the 3 different *trans*-regulatory groups it holds that *w*_1_ > *w*_2_ > *w*_3_. Therefore, in **(a)**, the assumption that differences in fitness correspond to differences in *s*_het_ is satisfied. In **(b)**, the assumption is violated.

**Supplementary Figure S3.**
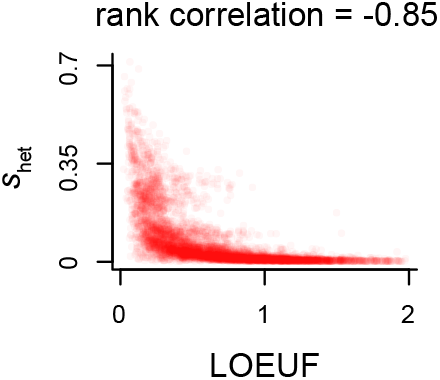
Assessing the concordance between two different measures of selective pressure against heterozygous coding loss-of-function alleles. Scatterplot of LOEUF estimates from gnomAD against shet estimates from Cassa et al. (2017). Each point corresponds to a gene.

**Supplementary Figure S4.**
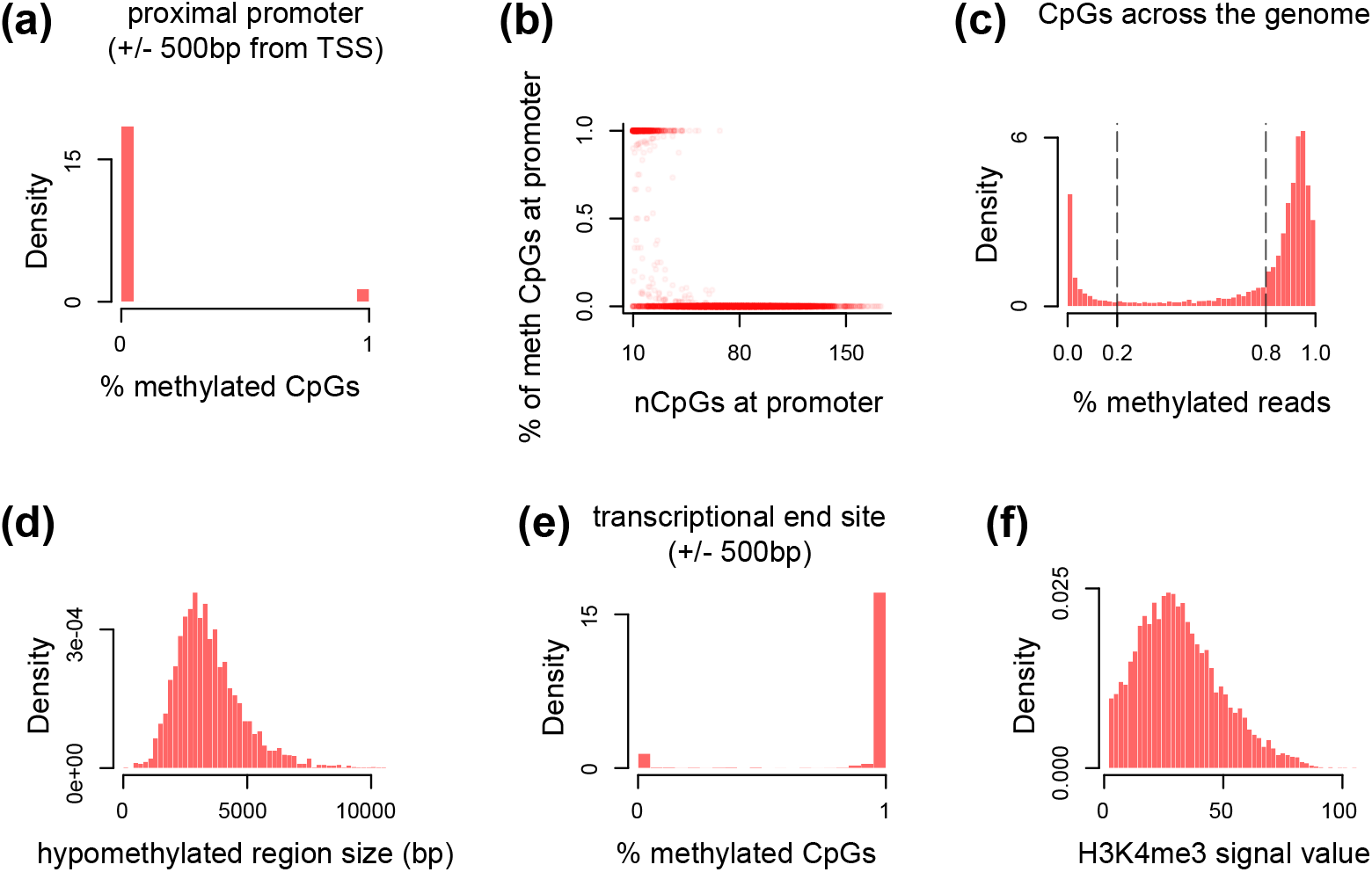
Variation exhibited by different epigenetic marks across the genome. **(a)** The distribution of DNA methylation across proximal promoters in the male germline. **(b)** The relationship between the total number of CpGs and the percentage of methylated CpGs across proximal promoters in the male germline. Each point corresponds to one of the 7,235 promoters used in our analysis. **(c)** The distribution of the percentage of sequencing reads supporting the methylated state at the individual CpG level, shown for all human CpGs with greater than 10x coverage (26,365,461 CpGs in total). The dashed vertical lines correspond to the thresholds of 0.2 and 0.8 that we used for labeling individual CpGs as hypomethylated and methylated, respectively. **(d)** The distribution of the size of the hypomethylated region at genes whose proximal promoter is hypomethylated in the male germline. **(e)** The distribution of DNA methylation at the transcriptional end site in the male germline. **(f)** The distribution of the intensity (height) of the H3K4me3 ChIP-seq signal at genes whose proximal promoter has at least one H3K4me3 peak in embryonic stem cells.

**Supplementary Figure S5.**
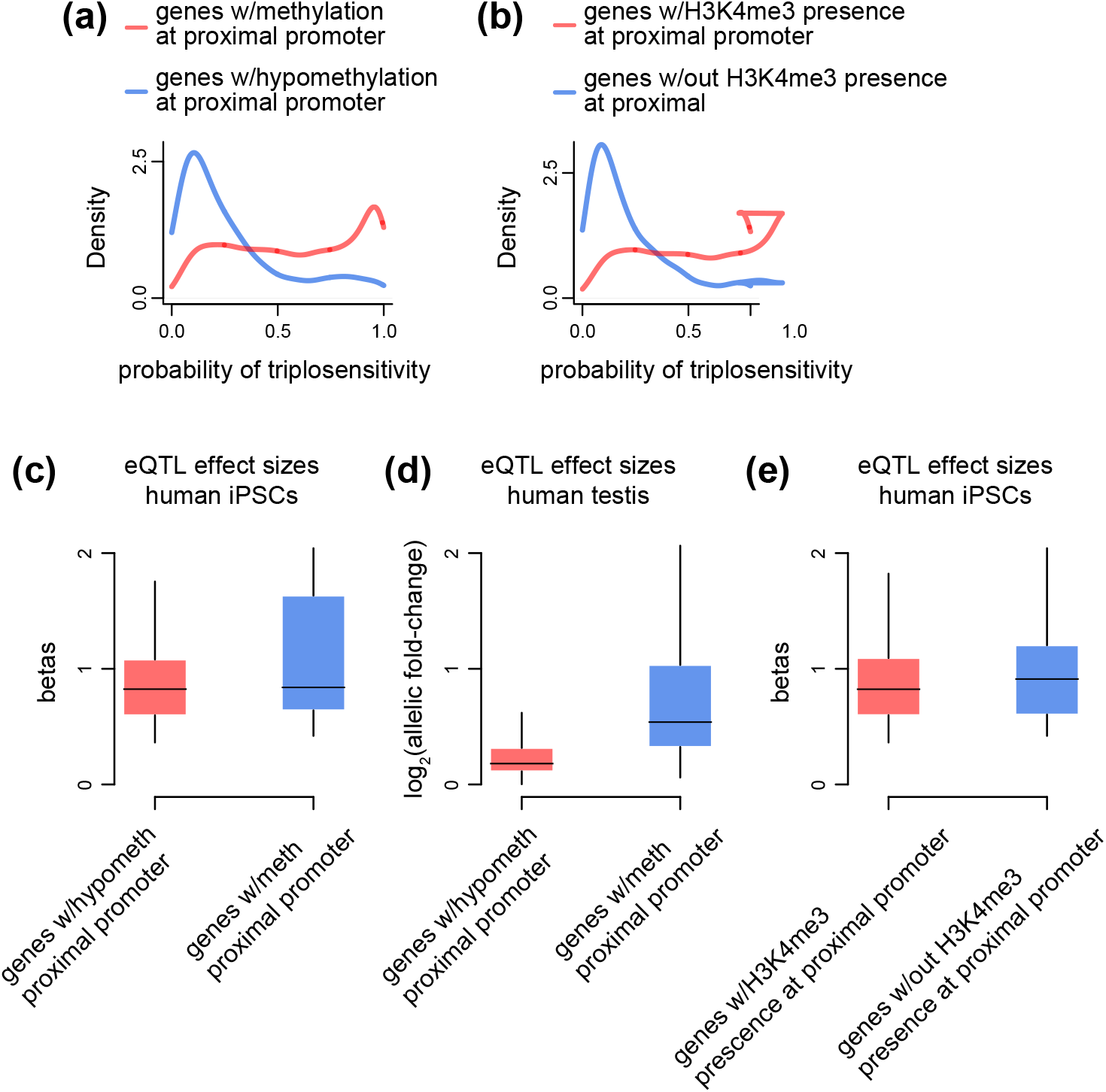
The relationship between genic intolerance to expression level elevation and the presence of proximal promoter DNA methylation/H3K4me3 peaks. **(a)** The distribution of genic triplosensitivity (Methods), for genes with methylated proximal promoters compared to genes with hypomethylated proximal promoters. **(b)** Like (a), but for genes with at least one H3K4me3 peak in the proximal promoter versus genes without any H3K4me3 peak in the proximal promoter. **(c)** The distribution of eQTL effect sizes (betas), estimated from human induced pluripotent stem cells (iPSCs), stratified according to proximal promoter methylation status in the male germline (Methods). **(d)** Like (c), but for testis eQTLs, where the effect size is represented by the log_2_ of allelic fold change following standard GTEx practices (Methods). **(e)** Like (c), but with genes stratified according to whether the proximal promoter has at least one H3K4me3 peak in h1-ESCs or not. For **(c-e)**, only eQTLs with q-value less than 0.05 and positive effect size (in log_2_ scale) are analyzed.

**Supplementary Figure S6.**
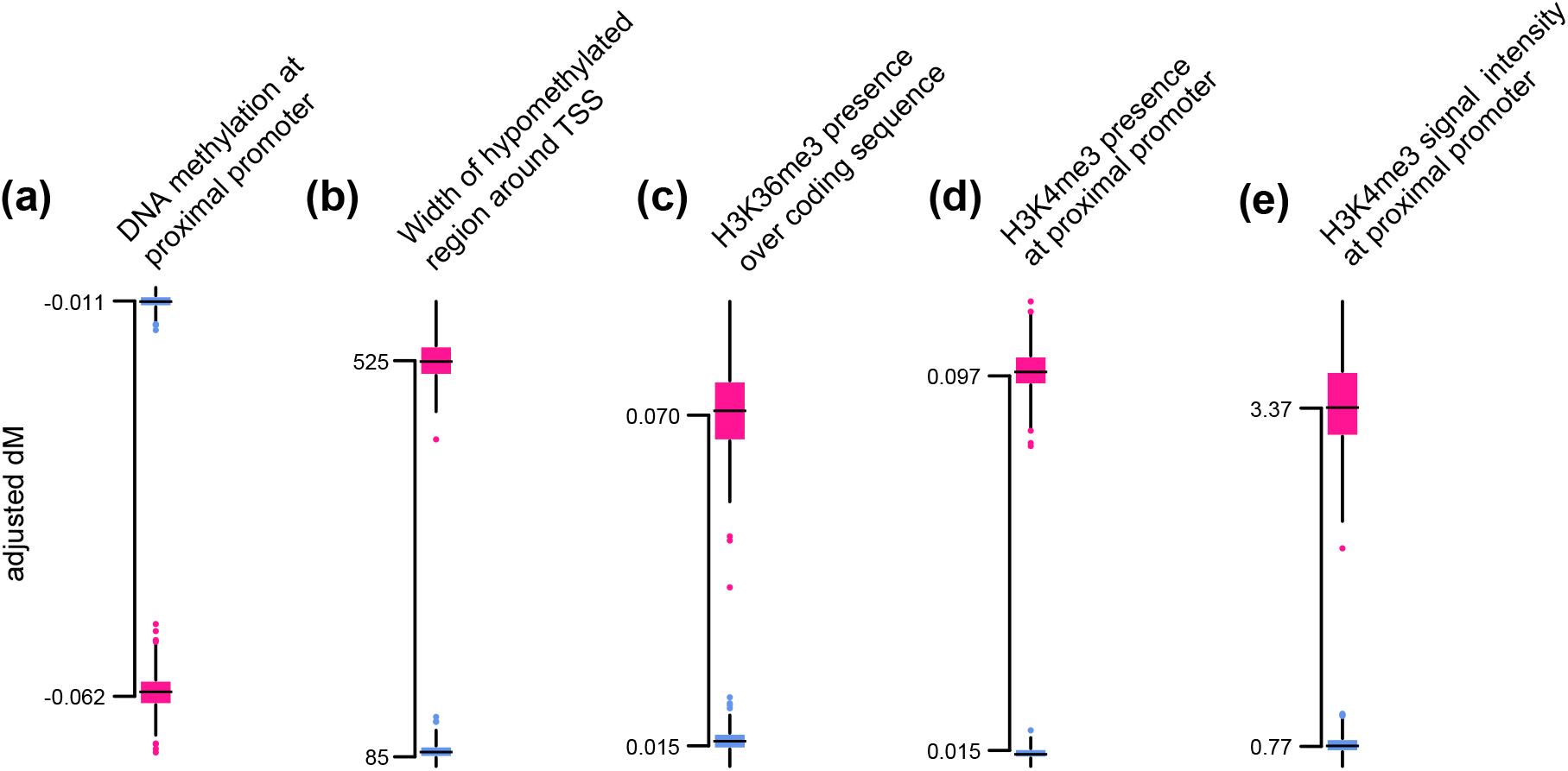
Testing whether the selective pressure on different epigenetic marks is a passive consequence of selection on gene expression in fetal cell types. **(a)** - **(e)** For different epigenetic marks, we computed the adjusted *dM* statistic and its corresponding null distribution for each of 172 fetal cell types (see Methods). We depict the distribution of adjusted *dM* (pink, 172 points) and compare it to the distribution of the most extreme null permutation (blue); maximum if the epigenetic mark is under positive selection, minimum if it is under negative selection.

**Supplementary Figure S7.**
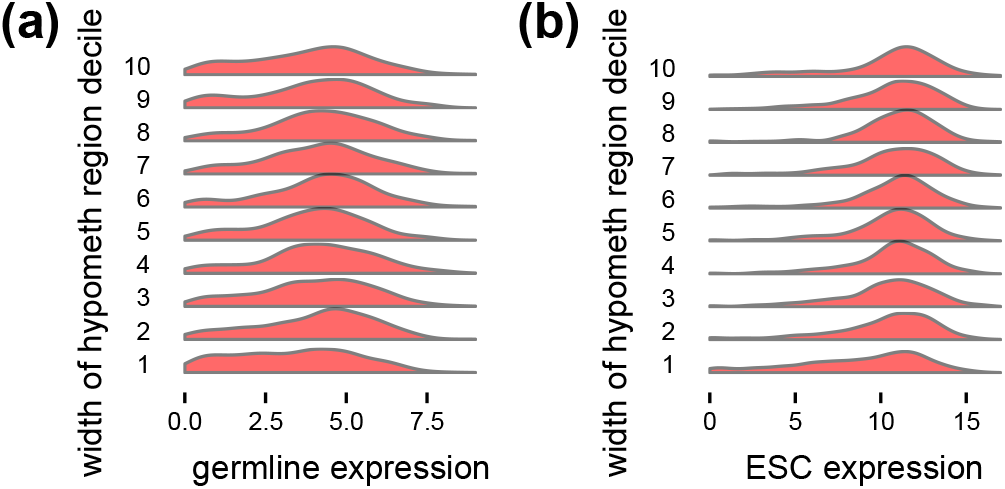
Assessing whether the positive selection on the size of the hypomethylated region around hypomethylated proximal promoters in the germline is explained by a causal role in gene regulation. The distribution of gene expression levels in testis (**(a)**) or embryonic stem cells (**(b)**), stratified according to the width of the hypomethylated region around the proximal promoter in the germline. Only genes with hypomethylated proximal promoters are analyzed.

**Supplementary Figure S8.**
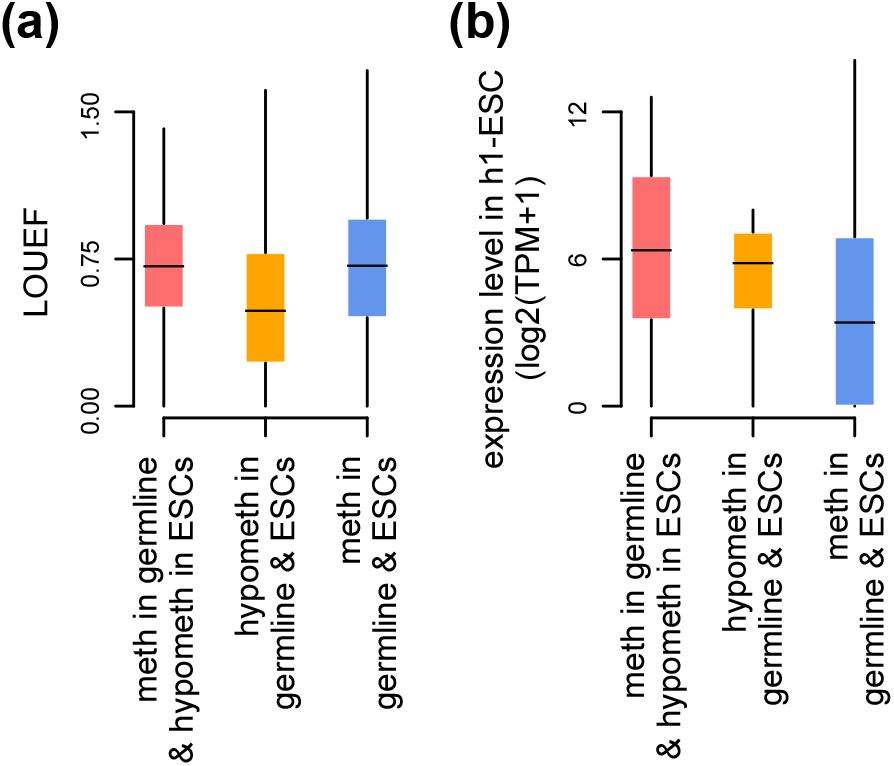
Assessing whether a causal role of proximal promoter DNA methylation methylation on expression during early pre-implantation development completely accounts for the inferred selective pressure. **(a)** The distribution of LOEUF estimates, stratified according to proximal promoter methylation status in the germline and embryonic stem cells (see Methods for details). **(b)** The distribution of expression level in embryonic stem cells, stratified according to proximal promoter methylation status in the germline and embryonic stem cells. The same genes as in (a).

**Supplementary Figure S9.**
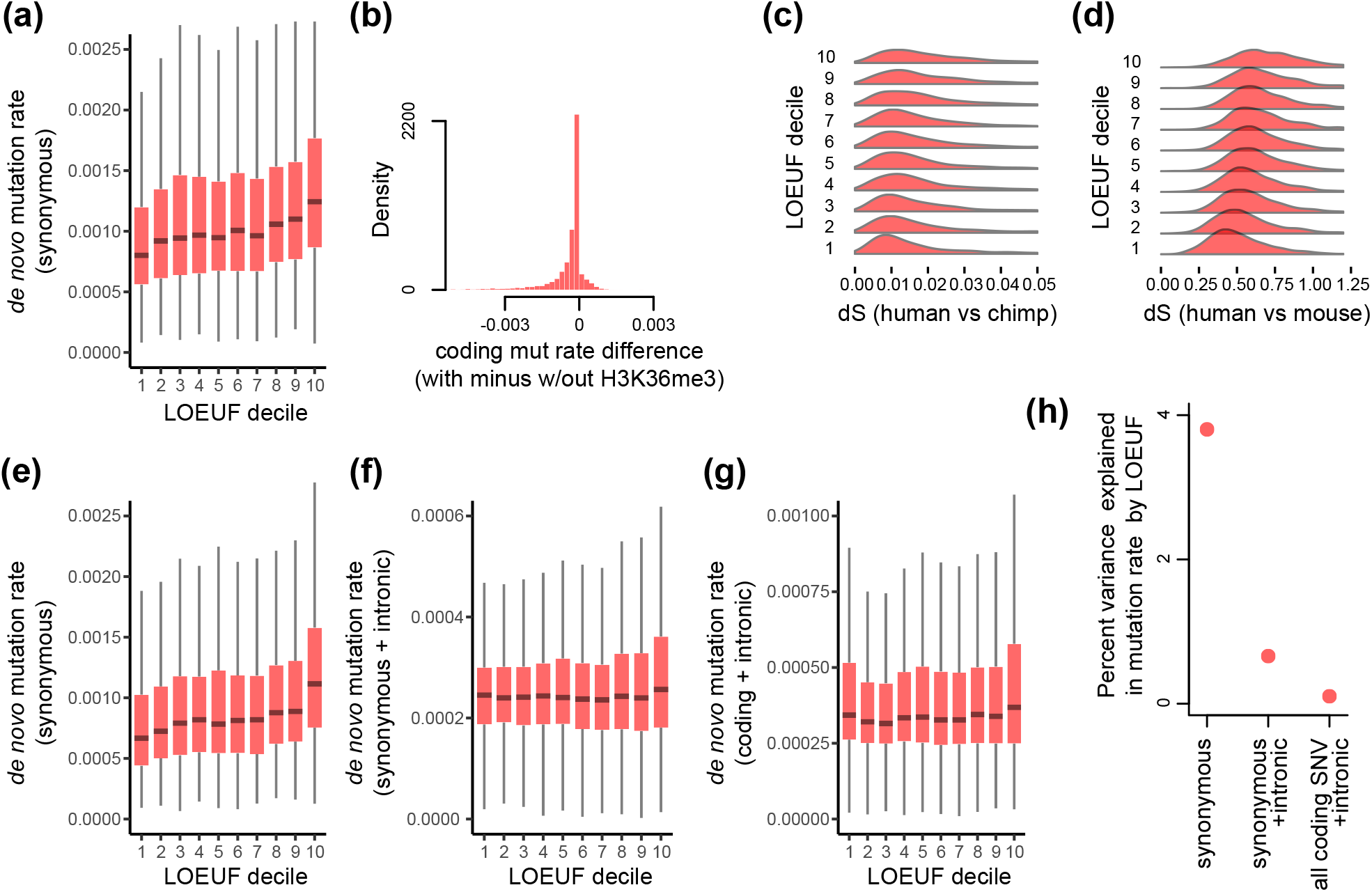
Coding *de novo* mutation rate, loss-of-function intolerance, and coding H3K36me3 patterns; additional analyses. **(a)** and **(b)**. Like Figure 4a and b, respectively, but where the number of synonymous mutations is computed by also adding a pseudocount of 1 to each gene. **(c)** and **(d)**. The distribution of the synonymous substitution rate (dS; obtained from Ensembl version 96 for human vs chimp and human vs mouse), stratified according to loss-of-function intolerance. **(e)** - **(g)**. Like Figure 4a, but with the mutation rate calculated using different types of *de novo* mutations (y axis label). **(h)** Percentage of variance in genic mutation rate explained by loss-of-function intolerance (LOEUF), for different types of *de novo* mutations used to calculate the mutation rate.

**Supplementary Figure S10.**
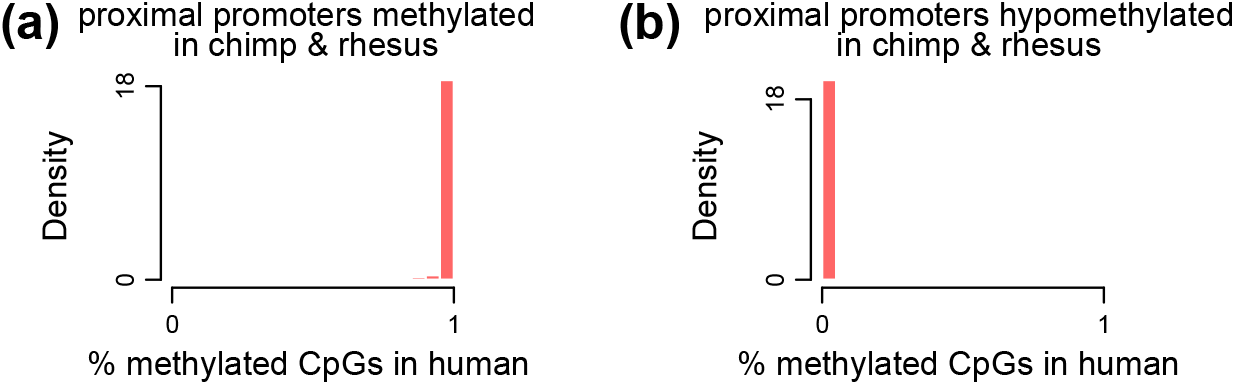
Comparative analysis of proximal promoter methylation. Methylation state in the human male germline of proximal promoters methylated (**a**) or hypomethylated (**b**) in chimp and rhesus male germline. In this analysis, rhesus serves as an outgroup.

**Supplementary Figure S11.**
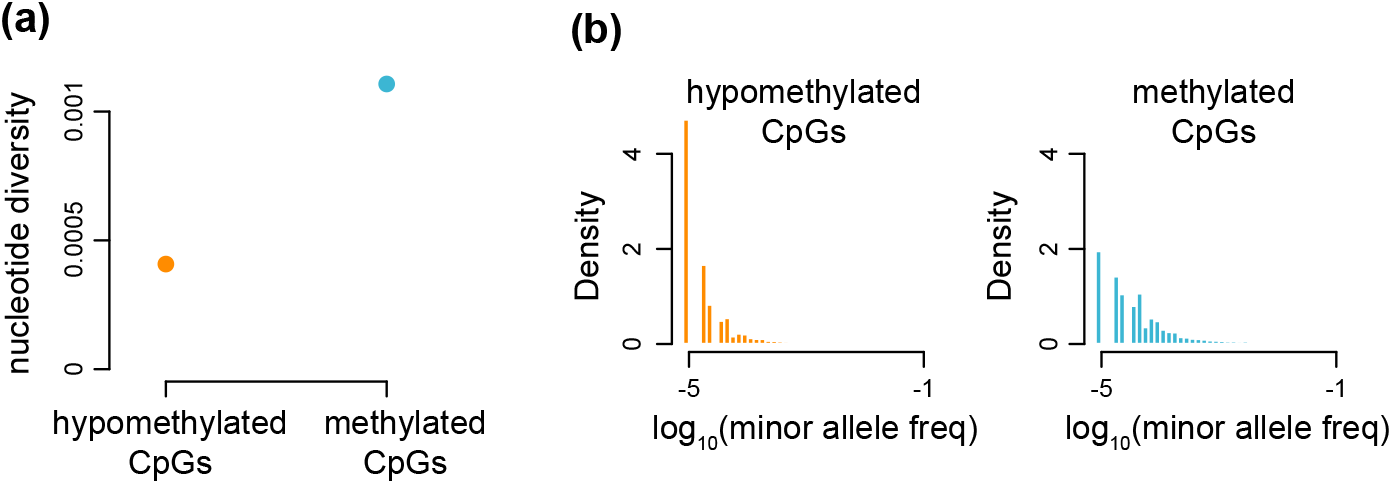
Genetic diversity of promoter CpG sites in TOPMed. **(a)** The nucleotide diversity of all promoter CpG sites, stratified according to whether they are methylated or hypomethylated in the male germline (≥ 80% and ≤20% of bisulfite sequencing reads supporting the methylated state, respectively). Nucleotide diversity was estimated as described in methods. Only CpGs with ≥ 10× coverage are considered. **(b)** The minor allele frequency spectrum of the CpGs used in (a).

## References

Antequera F, Bird A (2018). CpG Islands: A Historical Perspective. Methods in Molecular Biology 1766: 3–13. DOI:10.1007/978-1-4939-7768-0_1.

Asthana S, Noble WS, Kryukov G, Grant CE, Sunyaev S, Stamatoyannopoulos JA (2007). Widely distributed noncoding purifying selection in the human genome. PNAS 104.30: 12410–12415. DOI:10.1073/pnas.0705140104.

Bamshad M, Wooding SP (2003). Signatures of natural selection in the human genome. Nature Reviews Genetics 4.2: 99–111. DOI:10.1038/nrg999.

Bhattacharya S, Workman JL (2020). Regulation of SETD2 stability is important for the fidelity of H3K36me3 deposition. Epigenetics & Chromatin 13.1: 40. DOI:10.1186/s13072-020-00362-8.

Bintu L, Yong J, Antebi YE, McCue K, Kazuki Y, Uno N, Oshimura M, Elowitz MB (2016). Dynamics of epigenetic regulation at the single-cell level. Science 351.6274: 720–724. DOI:10.1126/science.aab2956.

Bird AP (1984). DNA methylation versus gene expression. Journal of Embryology and Experimental Morphology 83 Suppl: 31–40.

Blekhman R, Man O, Herrmann L, Boyko AR, Indap A, Kosiol C, Bustamante CD, Teshima KM, Przeworski M (2008). Natural selection on genes that underlie human disease susceptibility. Current Biology 18.12: 883–889. DOI:10.1016/j.cub.2008.04.074.

Boukas L, Bjornsson HT, Hansen KD (2020a). Promoter CpG Density Predicts Downstream Gene Loss-of-Function Intolerance. American Journal of Human Genetics 107.3: 487–498. DOI:10.1016/j.ajhg.2020.07.014.

Boukas L, Bjornsson HT, Hansen KD (2020b). Purifying selection acts on germline methylation to modify the CpG mutation rate at promoters. bioRxiv: 2020.07.04.187880. DOI:10.1101/2020.07.04.187880.

Boukas L, Havrilla JM, Hickey PF, Quinlan AR, Bjornsson HT, Hansen KD (2019). Coexpression patterns define epigenetic regulators associated with neurological dysfunction. Genome Research 29.4: 532–542. DOI:10.1101/gr.239442.118.

Boulard M, Edwards JR, Bestor TH (2015). FBXL10 protects Polycomb-bound genes from hypermethylation. Nature Genetics 47.5: 479–485. DOI:10.1038/ng.3272.

Cao J, O’Day DR, Pliner HA, Kingsley PD, Deng M, Daza RM, Zager MA, Aldinger KA, Blecher-Gonen R, Zhang F, et al. (2020). A human cell atlas of fetal gene expression. Science 370.6518. DOI:10.1126/science.aba7721.

Cassa CA, Weghorn D, Balick DJ, Jordan DM, Nusinow D, Samocha KE, O’Donnell-Luria A, MacArthur DG, Daly MJ, Beier DR, et al. (2017). Estimating the selective effects of heterozygous protein-truncating variants from human exome data. Nature Genetics 49.5: 806–810. DOI:10.1038/ng.3831.

Clouaire T, Webb S, Skene P, Illingworth R, Kerr A, Andrews R, Lee JH, Skalnik D, Bird A (2012). Cfp1 integrates both CpG content and gene activity for accurate H3K4me3 deposition in embryonic stem cells. Genes & Development 26.15: 1714–1728. DOI:10.1101/gad.194209.112.

Cohen NM, Kenigsberg E, Tanay A (2011). Primate CpG islands are maintained by heterogeneous evolutionary regimes involving minimal selection. Cell 145.5: 773–786. DOI:10.1016/j.cell.2011.04.024.

Collins RL, Glessner JT, Porcu E, Niestroj LM, Ulirsch J, Kellaris G, Howrigan DP, Everett S, Mohajeri K, Nuttle X, et al. (2021). A cross-disorder dosage sensitivity map of the human genome. bioRxiv: 2021.01.26.21250098. DOI:10.1101/2021.01.26.21250098.

Cooper DN (1983). Eukaryotic DNA methylation. Human Genetics 64.4: 315–333. DOI:10.1007/BF00292363.

Cooper DN, Youssoufian H (1988). The CpG dinucleotide and human genetic disease. Human Genetics 78.2: 151–155. DOI:10.1007/BF00278187.

Coulondre C, Miller JH, Farabaugh PJ, Gilbert W (1978). Molecular basis of base substitution hotspots in Escherichia coli. Nature 274.5673: 775–780. DOI:10.1038/274775a0.

Crow JF, Kimura M (2009). An introduction to population genetics theory. Blackburn Press.

DeBoever C, Li H, Jakubosky D, Benaglio P, Reyna J, Olson KM, Huang H, Biggs W, Sandoval E, D’Antonio M, et al. (2017). Large-Scale Profiling Reveals the Influence of Genetic Variation on Gene Expression in Human Induced Pluripotent Stem Cells. Cell Stem Cell 20.4: 533–546.e7. DOI:10.1016/j.stem.2017.03.009.

ENCODE Project Consortium (2012). An integrated encyclopedia of DNA elements in the human genome. Nature 489.7414: 57–74. DOI:10.1038/nature11247.

Eres IE, Gilad Y (2021). A TAD Skeptic: Is 3D Genome Topology Conserved? Trends in Genetics 37.3: 216–223. DOI:10.1016/j.tig.2020.10.009.

Falconer DS, MacKay TFC (1995). Introduction to Quantitative Genetics. 4th ed. London, England: Longman.

Finnegan AI, Kim S, Jin H, Gapinske M, Woods WS, Perez-Pinera P, Song JS (2020). Epigenetic engineering of yeast reveals dynamic molecular adaptation to methylation stress and genetic modulators of specific DNMT3 family members. Nucleic Acids Research 48.8: 4081–4099. DOI: 10.1093/nar/gkaa161.

Fisher RA (2015). The genetical theory of natural selection - scholar’s choice edition. Scholar’s Choice.

Frigo M, Johnson SG (2005). The design and implementation of FFTW3. Proceedings of the IEEE. Institute of Electrical and Electronics Engineers 93.2: 216–231. DOI:10.1109/jproc.2004.840301.

Fuller ZL, Berg JJ, Mostafavi H, Sella G, Przeworski M (2019). Measuring intolerance to mutation in human genetics. Nature Genetics 51.5: 772–776. DOI:10.1038/s41588-019-0383-1.

Gao Z, Moorjani P, Sasani TA, Pedersen BS, Quinlan AR, Jorde LB, Amster G, Przeworski M (2019). Overlooked roles of DNA damage and maternal age in generating human germline mutations. PNAS 116.19: 9491–9500. DOI:10.1073/pnas.1901259116.

Gardiner-Garden M, Frommer M (1987). CpG islands in vertebrate genomes. Journal of Molecular Biology 196.2: 261–282. DOI:10.1016/0022-2836(87)90689-9.

Gastonguay MS, Keele GR, Churchill GA (2022). “The Trouble with Triples: Examining the Impact of Measurement Error in Mediation Analysis”. DOI:10.1101/2022.07.07.499004.

Ginno PA, Gaidatzis D, Feldmann A, Hoerner L, Imanci D, Burger L, Zilbermann F, Peters AHFM, Edenhofer F, Smallwood SA, et al. (2020). A genome-scale map of DNA methylation turnover identifies site-specific dependencies of DNMT and TET activity. Nature Communications 11.1: 2680. DOI:10.1038/s41467-020-16354-x.

GTEx Consortium (2020). The GTEx Consortium atlas of genetic regulatory effects across human tissues. Science 369.6509: 1318–1330. DOI:10.1126/science.aaz1776.

GTEx Consortium, Laboratory, Data Analysis & Coordinating Center (LDACC)—Analysis Working Group, Statistical Methods groups—Analysis Working Group, Enhancing GTEx (eGTEx) groups, NIH Common Fund, NIH/NCI, NIH/NHGRI, NIH/NIMH, NIH/NIDA, Biospecimen Collection Source Site—NDRI, et al. (2017). Genetic effects on gene expression across human tissues. Nature 550.7675: 204–213. DOI:10.1038/nature24277.

Haigh J (1978). The accumulation of deleterious genes in a population–Muller’s Ratchet. Theoretical Population Niology 14.2: 251–267. DOI:10.1016/0040-5809(78)90027-8.

Hartl D, Krebs AR, Grand RS, Baubec T, Isbel L, Wirbelauer C, Burger L, Schübeler D (2019). CG dinucleotides enhance promoter activity independent of DNA methylation. Genome Research 29.4: 554–563. DOI:10.1101/gr.241653.118.

Heberle E, Bardet AF (2019). Sensitivity of transcription factors to DNA methylation. Essays Biochem. 63.6: 727–741.

Imbens GW, Rubin DB (2015). Causal Inference in Statistics, Social, and Biomedical Sciences. Cambridge University Press.

Irizarry RA, Ladd-Acosta C, Wen B, Wu Z, Montano C, Onyango P, Cui H, Gabo K, Rongione M, Webster M, et al. (2009). The human colon cancer methylome shows similar hypo- and hypermethylation at conserved tissue-specific CpG island shores. Nature Genetics 41.2: 178–186. DOI:10.1038/ng.298.

James LR, Mulaik SA, Brett JM (2006). A Tale of Two Methods.Organizational Research Methods 9.2: 233–244. DOI:10.1177/1094428105285144.

Jørgensen HF, Bird A (2002). MeCP2 and other methyl-CpG binding proteins. Ment Retard Dev Disabil Res Rev 8: 87–93. DOI:10.1002/mrdd.10021.

Karczewski KJ, Francioli LC, Tiao G, Cummings BB, Alföldi J, Wang Q, Collins RL, Laricchia KM, Ganna A, Birnbaum DP, et al. (2020). The mutational constraint spectrum quantified from variation in 141,456 humans. Nature 581.7809: 434–443. DOI:10.1038/s41586-020-2308-7.

Kimura M, Maruyama T (1966). The mutational load with epistatic gene interactions in fitness. Genetics 54.6: 1337–1351. DOI:10.1093/genetics/54.6.1337.

Kimura M (1967). On the evolutionary adjustment of spontaneous mutation rates*. Genetics Research 9.1: 23–34. DOI:10.1017/S0016672300010284.

Kondrashov AS (1995). Modifiers of mutation-selection balance: general approach and the evolution of mutation rates. Genetics Research 66.1: 53–69. DOI:10.1017/S001667230003439X.

Kong A, Frigge ML, Masson G, Besenbacher S, Sulem P, Magnusson G, Gudjonsson SA, Sigurdsson A, Jonasdottir A, Jonasdottir A, et al. (2012). Rate of de novo mutations and the importance of father’s age to disease risk. Nature 488.7412: 471–475. DOI:10.1038/nature11396.

Korthauer K, Irizarry RA (2018). Genome-wide repressive capacity of promoter DNA methylation is revealed through epigenomic manipulation. bioRxiv: 381145. DOI:10.1101/381145.

Krebs AR, Dessus-Babus S, Burger L, Schübeler D (2014). High-throughput engineering of a mammalian genome reveals building principles of methylation states at CG rich regions. eLife 3: e04094. DOI:10.7554/eLife.04094.

Kryazhimskiy S, Plotkin JB (2008). The population genetics of dN/dS. PLOS Genetics 4.12: e1000304. DOI:10.1371/journal.pgen.1000304.

Kryukov GV, Schmidt S, Sunyaev S (2005). Small fitness effect of mutations in highly conserved non-coding regions. Human Molecular Genetics 14.15: 2221–2229. DOI:10.1093/hmg/ddi226.

Landt SG, Marinov GK, Kundaje A, Kheradpour P, Pauli F, Batzoglou S, Bernstein BE, Bickel P, Brown JB, Cayting P, et al. (2012). ChIP-seq guidelines and practices of the ENCODE and modENCODE consortia. Genome Research 22.9: 1813–1831. DOI:10.1101/gr.136184.111.

Langmead B, Salzberg SL (2012). Fast gapped-read alignment with Bowtie 2. Nature Methods 9.4: 357–359. DOI:10.1038/nmeth.1923.

Lappalainen T, Greally JM (2017). Associating cellular epigenetic models with human phenotypes. Nature Reviews Genetics 18.7:441–451. DOI:10.1038/nrg.2017.32.

Ledgerwood A, Shrout PE (2011). The trade-off between accuracy and precision in latent variable models of mediation processes. J. Pers. Soc. Psychol. 101.6: 1174–1188. DOI:10.1037/a0024776.

Lee JH, Voo KS, Skalnik DG (2001). Identification and characterization of the DNA binding domain of CpG-binding protein. The Journal of Biological Chemistry 276.48: 44669–44676. DOI:10.1074/jbc.M107179200.

Leigh (1970). Natural Selection and Mutability. The American Naturalist 104.937: 301–305.

Lek M, Karczewski KJ, Minikel EV, Samocha KE, Banks E, Fennell T, O’Donnell-Luria AH, Ware JS, Hill AJ, Cummings BB, et al. (2016). Analysis of protein-coding genetic variation in 60,706 humans. Nature 536.7616: 285–291. DOI:10.1038/nature19057.

Li F, Mao G, Tong D, Huang J, Gu L, Yang W, Li GM (2013). The histone mark H3K36me3 regulates human DNA mismatch repair through its interaction with MutS*α*. Cell 153.3: 590–600. DOI:10.1016/j.cell.2013.03.025.

Liu H, Zhang J (2022). Is the Mutation Rate Lower in Genomic Regions of Stronger Selective Constraints? Molecular Biology and Evolution 39.8. DOI:10.1093/molbev/msac169.

Liu XS, Wu H, Ji X, Stelzer Y, Wu X, Czauderna S, Shu J, Dadon D, Young RA, Jaenisch R (2016). Editing DNA Methylation in the Mammalian Genome. Cell 167.1: 233–247.e17. DOI:10.1016/j.cell.2016.08.056.

Liu XS, Wu H, Krzisch M, Wu X, Graef J, Muffat J, Hnisz D, Li CH, Yuan B, Xu C, et al. (2018). Rescue of Fragile X Syndrome Neurons by DNA Methylation Editing of the FMR1 Gene. Cell 172.5: 979–992.e6. DOI:10.1016/j.cell.2018.01.012.

Lynch M (2007). The evolution of genetic networks by non-adaptive processes. Nature Reviews Genetics 8.10: 803–813. DOI:10.1038/nrg2192.

Lynch M (2011). The lower bound to the evolution of mutation rates. Genome Biology and Evolution 3: 1107–1118. DOI: 10.1093/gbe/evr066.

Lynch M, Ackerman MS, Gout JF, Long H, Sung W, Thomas WK, Foster PL (2016). Genetic drift, selection and the evolution of the mutation rate. Nature Reviews Genetics 17.11: 704–714. DOI:10.1038/nrg.2016.104.

Lynch M, Field MC, Goodson HV, Malik HS, Pereira-Leal JB, Roos DS, Turkewitz AP, Sazer S (2014). Evolutionary cell biology: two origins, one objective. PNAS 111.48: 16990–16994. DOI: 10.1073/pnas.1415861111.

Lynch M, Trickovic B (2020). A Theoretical Framework for Evolutionary Cell Biology. Journal of Molecular Biology 432.7: 1861–1879. DOI:10.1016/j.jmb.2020.02.006.

MacKinnon DP (2012). Introduction to statistical mediation analysis. Routledge.

Maia LP, Botelho DF, Fontanari JF (2003). Analytical solution of the evolution dynamics on a multiplicative-fitness landscape. Journal of Mathematical Biology 47.5: 453–456. DOI:10.1007/s00285-003-0208-8.

Marinov, Kundaje (2018). ChIP-ping the branches of the tree: functional genomics and the evolution of eukaryotic gene regulation. Briefings in Functional Genomics 17.2: 116–137.

Martincorena I, Seshasayee ASN, Luscombe NM (2012). Evidence of non-random mutation rates suggests an evolutionary risk management strategy. Nature 485.7396: 95–98. DOI:10.1038/nature10995.

Mendoza A de, Nguyen TV, Ford E, Poppe D, Buckberry S, Pflueger J, Grimmer MR, Stolzenburg S, Bogdanovic O, Oshlack A, et al. (2022). Large-scale manipulation of promoter DNA methylation reveals context-specific transcriptional responses and stability. Genome Biol. 23.1: 163.

Molaro A, Hodges E, Fang F, Song Q, McCombie WR, Hannon GJ, Smith AD (2011). Sperm methylation profiles reveal features of epigenetic inheritance and evolution in primates. Cell 146.6: 1029–1041. DOI:10.1016/j.cell.2011.08.016.

Monroe JG, Srikant T, Carbonell-Bejerano P, Becker C, Lensink M, Exposito-Alonso M, Klein M, Hildebrandt J, Neumann M, Kliebenstein D, et al. (2022). Mutation bias reflects natural selection in Arabidopsis thaliana. Nature 602.7895: 101–105. DOI:10.1038/s41586-021-04269-6.

Morselli M, Pastor WA, Montanini B, Nee K, Ferrari R, Fu K, Bonora G, Rubbi L, Clark AT, Ottonello S, et al. (2015). In vivo targeting of de novo DNA methylation by histone modifications in yeast and mouse. eLife 4: e06205. DOI:10.7554/eLife.06205.

Murray SC, Lorenz P, Howe FS, Wouters M, Brown T, Xi S, Fischl H, Khushaim W, Rayappu JR, Angel A, et al. (2019). H3K4me3 is neither instructive for, nor informed by, transcription. bioRxiv:709014. DOI:10.1101/709014.

Musselman CA, Lalonde ME, Côté J, Kutateladze TG (2012). Perceiving the epigenetic landscape through histone readers. Nat Struct Mol Biol 19: 1218–1227. DOI:10.1038/nsmb.2436.

Nielsen R (2005). Molecular signatures of natural selection. Annual Review of Genetics 39: 197–218. DOI:10.1146/annurev.genet.39.073003.112420.

Nielsen R, Bustamante C, Clark AG, Glanowski S, Sackton TB, Hubisz MJ, Fledel-Alon A, Tanenbaum DM, Civello D, White TJ, et al. (2005). A scan for positively selected genes in the genomes of humans and chimpanzees. PLOS Biology 3.6: e170. DOI:10.1371/journal.pbio.0030170.

Nielsen R, Hellmann I, Hubisz M, Bustamante C, Clark AG (2007). Recent and ongoing selection in the human genome. Nature Reviews Genetics 8.11: 857–868. DOI:10.1038/nrg2187.

Nielsen R, Yang Z (2003). Estimating the distribution of selection coefficients from phylogenetic data with applications to mitochondrial and viral DNA. Molecular Biology and Evolution 20.8: 1231–1239. DOI:10.1093/molbev/msg147.

Ooi SKT, Qiu C, Bernstein E, Li K, Jia D, Yang Z, Erdjument-Bromage H, Tempst P, Lin SP, Allis CD, et al. (2007). DNMT3L connects unmethylated lysine 4 of histone H3 to de novo methylation of DNA. Nature 448.7154: 714–717. DOI:10.1038/nature05987.

Otter T, Pachali MJ, Mayer S, Landwehr JR (2018). Causal Inference Using Mediation Analysis or Instrumental Variables–Full Mediation in the Absence of Conditional Independence. Marketing: ZFP–Journal of Research and Management 40.2: 41–57. DOI:10.2139/ssrn.3135313.

Panchin AY, Makeev VJ, Medvedeva YA (2016). Preservation of methylated CpG dinucleotides in human CpG islands. Biology Direct 11.1: 11. DOI:10.1186/s13062-016-0113-x.

Patro R, Duggal G, Love MI, Irizarry RA, Kingsford C (2017). Salmon provides fast and bias-aware quantification of transcript expression. Nature Methods 14.4: 417–419. DOI:10.1038/nmeth.419.

Pearl J (2014). Interpretation and identification of causal mediation. Psychological Methods 19.4: 459–481. DOI:10.1037/a0036434.

Petrovski S, Wang Q, Heinzen EL, Allen AS, Goldstein DB (2013). Genic intolerance to functional variation and the interpretation of personal genomes. PLOS Genetics 9.8: e1003709. DOI:10.1371/journal.pgen.1003709.

Pfister SX, Ahrabi S, Zalmas LP, Sarkar S, Aymard F, Bachrati CZ, Helleday T, Legube G, La Thangue NB, Porter ACG, et al. (2014). SETD2-dependent histone H3K36 trimethylation is required for homologous recombination repair and genome stability. Cell Reports 7.6: 2006–2018. DOI:10.1016/j.celrep.2014.05.026.

Phipson B, Smyth GK (2010). Permutation P-values should never be zero: calculating exact P-values when permutations are randomly drawn. Statistical Applications in Genetics and Molecular Biology 9: Article39. DOI:10.2202/1544-6115.1585.

Pollard KS, Hubisz MJ, Rosenbloom KR, Siepel A (2010). Detection of nonneutral substitution rates on mammalian phylogenies. Genome Research 20.1: 110–121. DOI:10.1101/gr.097857.109.

Ptashne (2007). On the use of the word ‘epigenetic’. Current Biology 17.7: R233–R236. DOI:10.1016/j.cub.2007.02.030.

Qu J, Hodges E, Molaro A, Gagneux P, Dean MD, Hannon GJ, Smith AD (2018). Evolutionary expansion of DNA hypomethylation in the mammalian germline genome. Genome Research 28.2: 145–158. DOI:10.1101/gr.225896.117.

Quinlan AR, Hall IM (2010). BEDTools: a flexible suite of utilities for comparing genomic features. Bioinformatics 26.6: 841–842. DOI:10.1093/bioinformatics/btq033.

Rahim K (2021). *fftwtools: Wrapper for ‘FFTW3’ Includes: One-Dimensional, Two-Dimensional, ThreeDimensional, and Multivariate Transforms*. R package version 0.9-11.

Renciuk D, Blacque O, Vorlickova M, Spingler B (2013). Crystal structures of B-DNA dodecamer containing the epigenetic modifications 5-hydroxymethylcytosine or 5-methylcytosine. Nucleic Acids Research 41.21: 9891–9900. DOI:10.1093/nar/gkt738.

Rodriguez-Galindo M, Casillas S, Weghorn D, Barbadilla A (2020). Germline de novo mutation rates on exons versus introns in humans. Nature Communications 11.1: 3304. DOI:10.1038/s41467-020-17162-z.

Sabeti PC, Schaffner SF, Fry B, Lohmueller J, Varilly P, Shamovsky O, Palma A, Mikkelsen TS, Altshuler D, Lander ES (2006). Positive natural selection in the human lineage. Science 312.5780: 1614–1620. DOI:10.1126/science.1124309.

Soneson C, Love MI, Robinson MD (2015). Differential analyses for RNA-seq: transcript-level estimates improve gene-level inferences. F1000Research 4: 1521. DOI:10.12688/f1000research.7563.2.

Song Q, Decato B, Hong EE, Zhou M, Fang F, Qu J, Garvin T, Kessler M, Zhou J, Smith AD (2013). A reference methylome database and analysis pipeline to facilitate integrative and comparative epigenomics. PLOS One 8.12: e81148. DOI:10.1371/journal.pone.0081148.

Stadler MB, Murr R, Burger L, Ivanek R, Lienert F, Schöler A, Nimwegen E van, Wirbelauer C, Oakeley EJ, Gaidatzis D, et al. (2011). DNA-binding factors shape the mouse methylome at distal regulatory regions. Nature 480.7378: 490–495. DOI:10.1038/nature10716.

Sturtevant AH (1937). Essays on evolution. I. on the effects of selection on mutation rate. The Quarterly Review of Biology 12.4: 464–467. DOI:10.1086/394543.

Sun XJ, Wei J, Wu XY, Hu M, Wang L, Wang HH, Zhang QH, Chen SJ, Huang QH, Chen Z (2005). Identification and characterization of a novel human histone H3 lysine 36-specific methyltransferase. The Journal of Biological Chemistry 280.42: 35261–35271. DOI:10.1074/jbc.M504012200.

Sun Z, Zhang Y, Jia J, Fang Y, Tang Y, Wu H, Fang D (2020). H3K36me3, message from chromatin to DNA damage repair. Cell & Bioscience 10: 9. DOI:10.1186/s13578-020-0374-z.

Taliun D, Harris DN, Kessler MD, Carlson J, Szpiech ZA, Torres R, Taliun SAG, Corvelo A, Gogarten SM, Kang HM, et al. (2021). Sequencing of 53,831 diverse genomes from the NHLBI TOPMed Program. Nature 590.7845: 290–299. DOI:10.1038/s41586-021-03205-y.

Thomson JP, Skene PJ, Selfridge J, Clouaire T, Guy J, Webb S, Kerr ARW, Deaton A, Andrews R, James KD, et al. (2010). CpG islands influence chromatin structure via the CpG-binding protein Cfp1. Nature 464.7291: 1082–1086. DOI:10.1038/nature08924.

Tsuruta M, Sugitani Y, Sugimoto N, Miyoshi D (2021). Combined Effects of Methylated Cytosine and Molecular Crowding on the Thermodynamic Stability of DNA Duplexes. International Journal of Molecular Sciences 22.2. DOI:10.3390/ijms22020947.

VanderWeele TJ (2016). Mediation Analysis: A Practitioner’s Guide. Annual Review of Public Health 37: 17–32. DOI:10.1146/annurev-publhealth-032315-021402.

Wang RY, Kuo KC, Gehrke CW, Huang LH, Ehrlich M (1982). Heat- and alkali-induced deamination of 5-methylcytosine and cytosine residues in DNA. Biochimica et Biophysica Acta 697.3: 371–377. DOI:10.1016/0167-4781(82)90101-4.

Wang S, Kool ET (1995). Origins of the large differences in stability of DNA and RNA helices: C-5 methyl and 2’-hydroxyl effects. Biochemistry 34.12: 4125–4132. DOI:10.1021/bi00012a031.

Wang Z, Chivu AG, Choate LA, Rice EJ, Miller DC, Chu T, Chou SP, Kingsley NB, Petersen JL, Finno CJ, et al. (2022). Prediction of histone post-translational modification patterns based on nascent transcription data. Nature Genetics 54.3: 295–305. DOI:10.1038/s41588-022-01026-x.

Xia B, Yan Y, Baron M, Wagner F, Barkley D, Chiodin M, Kim SY, Keefe DL, Alukal JP, Boeke JD, et al. (2020). Widespread Transcriptional Scanning in the Testis Modulates Gene Evolution Rates. Cell 180.2: 248–262.e21. DOI:10.1016/j.cell.2019.12.015.

Xia B, Yanai I (2022). Gene expression levels modulate germline mutation rates through the compound effects of transcription-coupled repair and damage. Human Genetics 141.6: 1211–1222. DOI:10.1007/s00439-021-02355-3.

Yin Y, Morgunova E, Jolma A, Kaasinen E, Sahu B, Khund-Sayeed S, Das PK, Kivioja T, Dave K, Zhong F, et al. (2017). Impact of cytosine methylation on DNA binding specificities of human transcription factors. Science 356.6337. DOI:10.1126/science.aaj2239.

Yu DH, Ware C, Waterland RA, Zhang J, Chen MH, Gadkari M, Kunde-Ramamoorthy G, Nosavanh LM, Shen L (2013). Developmentally programmed 3’ CpG island methylation confers tissue-and cell-type-specific transcriptional activation. Molecular and Cellular Biology 33.9: 1845–1858. DOI:10.1128/MCB.01124-12.

Zhang Y, Liu T, Meyer CA, Eeckhoute J, Johnson DS, Bernstein BE, Nusbaum C, Myers RM, Brown M, Li W, et al. (2008). Model-based analysis of ChIP-Seq (MACS). Genome Biology 9.9: R137. DOI:10.1186/gb-2008-9-9-r137.

Zhao G, Li K, Li B, Wang Z, Fang Z, Wang X, Zhang Y, Luo T, Zhou Q, Wang L, et al. (2020). Gene4Denovo: an integrated database and analytic platform for de novo mutations in humans. Nucleic Acids Research 48.D1: D913–D926. DOI:10.1093/nar/gkz923.

Zhou Y, He F, Pu W, Gu X, Wang J, Su Z (2020). The Impact of DNA Methylation Dynamics on the Mutation Rate During Human Germline Development. G3 10.9: 3337–3346. DOI:10.1534/g3.120.401511.

